# RNA Transcripts Serve as a Template for Double-Strand Break Repair in Human Cells

**DOI:** 10.1101/2025.02.23.639725

**Authors:** Manisha Jalan, Alessandra Brambati, Hina Shah, Niamh McDermott, Juber Patel, Yingjie Zhu, Ahmet Doymaz, Julius Wu, Kyrie S Anderson, Andrea Gazzo, Fresia Pareja, Takafumi N Yamaguchi, Theodore Vougiouklakis, Sana Ahmed-Seghir, Philippa Steinberg, Anna Neiman-Golden, Benura Azeroglu, Joan Gomez-Aguilar, Edaise M da Silva, Suleman Hussain, Daniel Higginson, Paul C Boutros, Nadeem Riaz, Jorge S Reis-Filho, Simon N Powell, Agnel Sfeir

## Abstract

Double-strand breaks (DSBs) are toxic lesions that lead to genome instability. While canonical DSB repair pathways typically operate independently of RNA, emerging evidence suggests that RNA:DNA hybrids and transcripts near damaged sites can influence repair outcomes. However, a direct role for transcript RNA as a template during DSB repair in human cells is yet to be established. In this study, we designed fluorescent- and sequencing-based assays, which demonstrated that RNA-containing oligonucleotides and messenger RNA serve as templates to promote DSB repair. We conducted a CRISPR/Cas9-based genetic screen to identify factors that promote RNA-templated DSB repair (RT-DSBR), and of the candidate polymerases, we identified DNA polymerase-zeta (Polζ) as the potential reverse transcriptase that facilitates RT-DSBR. Furthermore, by analyzing sequencing data from cancer genomes, we identified the presence of whole intron deletions, a unique genomic scar reflective of RT-DSBR activity generated when spliced mRNA serves as the repair template. These findings highlight RT-DSBR as an alternative pathway for repairing DSBs in transcribed genes, with potential mutagenic consequences.

## INTRODUCTION

The human genome is constantly exposed to endogenous and exogenous insults that cause DNA damage. Among the various types of DNA damage, double-strand breaks (DSBs) are particularly harmful, leading to genome instability, a hallmark of aging, cancer, and neurodegeneration^1^. DSBs are repaired through three major pathways: homologous recombination (HR), non-homologous end-joining (NHEJ), and microhomology-mediated end-joining (MMEJ). NHEJ repairs DSBs by ligating broken ends with minimal processing. HR and MMEJ rely on DNA resection to generate a single-stranded DNA tail that anneals to the sister chromatid or the opposing DSB end^2^. While these canonical repair pathways generally function independently of RNA, more than 80% of the genome is actively transcribed at any given time^3^. As a result, DSB repair frequently occurs in open chromatin regions, and RNA transcription must be intricately coordinated with DNA repair to ensure genomic stability and appropriate gene expression.

Over the years, research into the interplay between transcription and DSB repair uncovered how DSBs modulate gene expression and how transcription, in turn, shapes repair outcomes. While the activation of DNA damage signaling kinases, ATM and DNA-PK, was shown to repress transcription by RNA Pol II near break sites^4–6^, conflicting results suggested that enhanced transcription generates noncoding RNAs that amplify DNA damage signaling and recruit HR factors to DSB sites^7–10^. RNA transcripts accumulating at break sites can anneal to DNA, leading to the formation of RNA:DNA hybrids^11–15^ that promote DSB repair by regulating DNA end resection and facilitating the recruitment of repair factors^16^. RNA transcripts have also been shown to stimulate HR by invading the donor DNA in response to DSBs, forming an intermediate D-loop containing RNA, which increases the accessibility of the break to the donor DNA template^17^.

Beyond its indirect role in orchestrating DSB repair, RNA can also play a more direct, instructive role by serving as a template for DSB repair. In *S. cerevisiae*, it has been demonstrated that messenger RNA (mRNA) can be reverse transcribed to act as a template for DSB repair in contexts where RNaseH1 and RNaseH2 are lacking. One form of RNA-templated DSR Repair (RT-DSBR) involves the production of a cDNA intermediate by Ty retrotransposons, which is then used as a template for HR repair. Another mechanism entails base pairing of the RNA with single-stranded DNA (ssDNA) flanking the break site, followed by its copying in cis by the translesion polymerase zeta (Polζ)^18–20^. Whether RT-DSBR is conserved in higher eukaryotes remains unknown.

In human cells, the transfer of genetic information from RNA to DNA is predominantly mediated by two reverse transcriptase activities. The first is telomerase, which reverse transcribes its RNA to replenish telomere DNA^21^. The second reverse transcriptase activity involves ORF2, which reverse transcribes LINE-1 RNA into DNA, allowing the integration of the transposable element into the genome^22^. Both ORF2 and telomerase activities have been detected at DSBs induced by CRISPR/Cas9 cleavage, where LINE-1 and TTAGGG are introduced, albeit with very low efficiency^23,24^. In addition, biochemical studies have shown that while several human replicative and translesion polymerases can copy up to 2-3 embedded ribonucleotides, Polymerase theta (Polθ-encoded by *POLQ*) can reverse transcribe several kilobases of mRNA *in vitro*^25^. This reverse transcriptase activity has been suggested to facilitate RNA-templated repair *in vivo*. However, whether mRNA can act as a template for DSB repair in human cells and which enzyme is responsible for mediating this reverse transcription remains uncertain.

Here, we investigated the direct role of RNA copying when templating DSB repair in human cells by developing complementary fluorescent and sequencing-based reporter assays. Our results demonstrate that RNA-containing oligonucleotides and RNA transcripts provide a donor sequence for RT-DSBR in human cells. Using CRISPR/Cas9 screening, we identified the translesion polymerase Polζ as the reverse transcriptase that promotes RT-DSBR. Furthermore, we predicted that by using a spliced mRNA as a template, RT-DSBR would lead to the deletion of an intron. We investigated the outcome of repairing a break within an intron using spliced transcript and demonstrated that this process results in the complete deletion of the intron from the genome. We leveraged the phenomenon of intron loss to provide evidence of RT-DSBR under physiological conditions. By analyzing sequencing datasets from MSK-IMPACT (Memorial Sloan Kettering-Integrated Mutation Profiling of Actionable Cancer Targets) and PCAWG (Pan-Cancer Analysis of Whole Genomes), we identified precise deletions of intronic sequences, which we refer to as whole intron deletions (WIDs). This analysis uncovered WID as a genomic signature indicative of RT-DSBR activity in cancer genomes, suggesting that spliced mRNA serves as a repair template for DSBs. Collectively, our findings suggest that RNA can serve as a template for repairing DSBs. We propose that RT-DSBR might be particularly significant in regions with high transcriptional activity by providing a mechanism for maintaining genome integrity.

## Results

### Human cells repair DSBs using RNA as a template to copy genetic information

To investigate whether RNA can directly serve as a template during DSB repair in human cells, we developed two complementary reporter assays capable of detecting reverse transcription activity at a CRISPR/Cas9-induced DSB using either fluorescence or sequencing readouts. In the first assay, we used a BFP (blue fluorescent protein)-to-GFP (green fluorescent protein) reporter system that involves a single amino acid change (His66Tyr) detectable by flow cytometry (Fig. 1A, S1A-B)^26^. As previously demonstrated, repairing a Cas9-induced DSB in a BFP gene, randomly integrated into the genome, successfully converted BFP to Green Fluorescent Protein (GFP). This conversion occurred when a single-stranded DNA donor (DNA^GFP^) containing the corresponding amino acid change was used as the repair template^27^. To adapt this reporter for RNA-templated DSB Repair (RT-DSBR), we generated chimeric oligonucleotide donors, replacing the three bases coding for tyrosine in the DNA template with the corresponding ribonucleotides (rNTPs), creating a series of donors with 3 to 15 ribonucleotides (DNA/RNA^3R/6R/8R/15R^) (Fig. 1B). The successful repair of BFP, resulting in GFP expression, is anticipated to be mediated by a reverse transcriptase that copies RNA residues into DNA.

**Figure 1.**
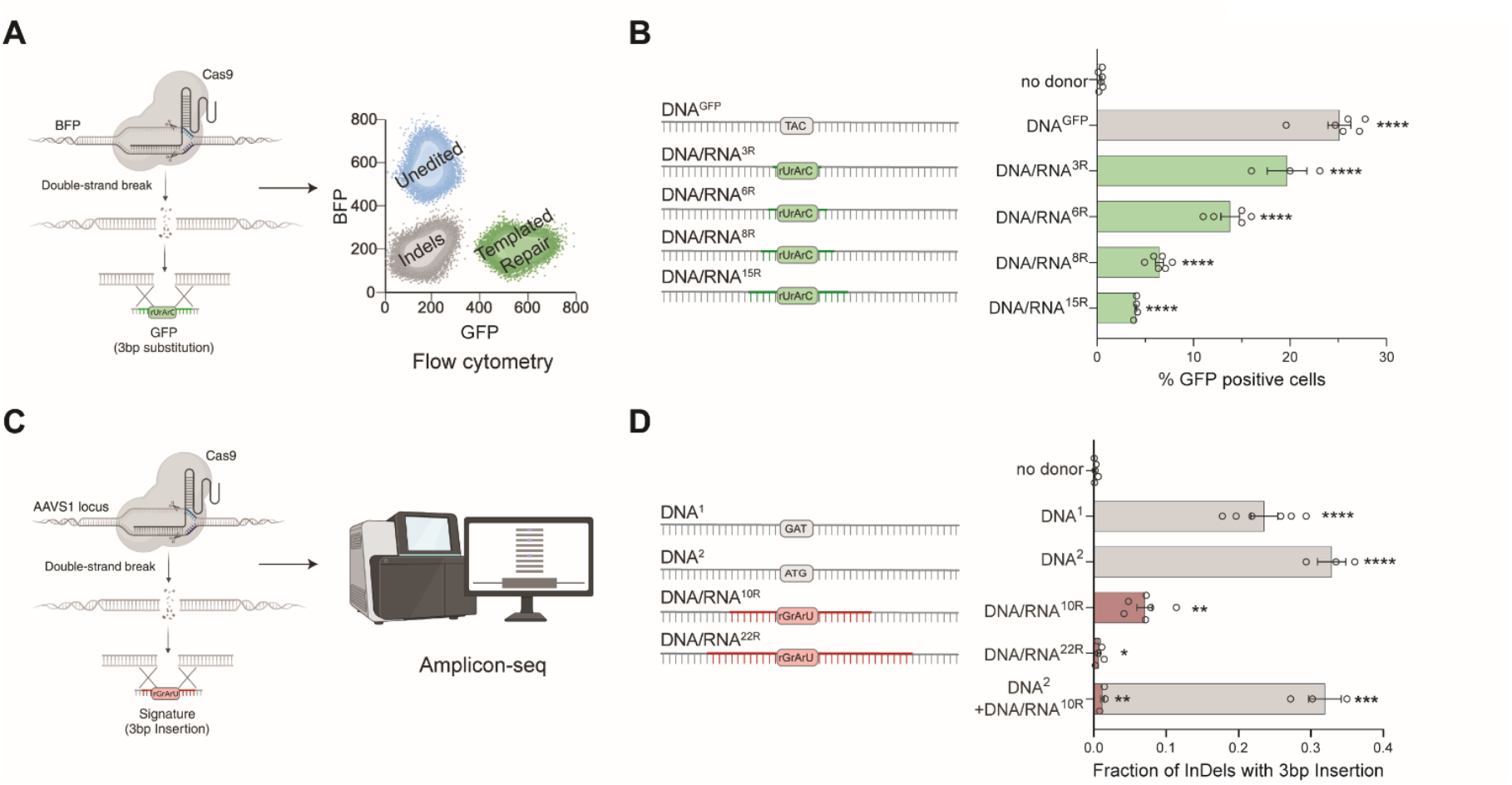
Human cells use RNA to template DSB repair. **A**, Schematic of the BFP-to-GFP assay designed to generate a green fluorescent signal via RNA templated DSB repair (RT-DSBR). This assay exploits the single amino acid change that differentiates Blue Fluorescent Protein (BFP) from Green Fluorescent Protein (GFP), switching the fluorescence from blue to green. A DSB is introduced at an integrated BFP locus using CRISPR/Cas9, and cells repair the break with a single-stranded DNA donor (DNA^GFP^) containing the GFP codon, switching from BFP to GFP fluorescence. To detect RT-DSBR activity, we used DNA/RNA chimeric donors where the sequence required to swap the codon was encoded by ribonucleotides instead of deoxyribonucleotides. **B**, Right: schematic of the 120bp chimeric donors used in the BFP-to-GFP assay with green segments representing stretches of ribonucleotides. Left: GFP signal quantification was performed by flow cytometry with different donors (n≥3) and compared to a non-donor control. **C**, Schematic of the AAVS1-seq assay. A targeted DSB is introduced at the AAVS1 genomic locus using CRISPR/Cas9, and the donor DNA or DNA/RNA chimeras containing a 3bp insertion are transfected into the cells. Successful repair using the donor leads to the incorporation of the mutational signature, which is detected by PCR amplification and Next Generation Sequencing. **D**, Right: a schematic of the 60 bp donor templates used in the AAVS1-seq assay, with red segments representing stretches of ribonucleotides. Left: quantification of the fraction of repair products containing the 3bp insertion signature after the Cas9 DSB is repaired by different donors, as measured by the AAVS1-seq assay (n=3) and compared to a non-donor control. For B and D: Statistical significance was assessed using unpaired Student’s t-test (* p < 0.05, ** p < 0.01, *** p < 0.001, ****p < 0.0001). Error bars represent the standard error of the mean (± SEM). See also Figure S1.

Using HEK293T cells stably expressing BFP, we delivered Cas9 protein and sgRNA targeting BFP, along with DNA/RNA chimera donors. In the presence of the DNA^GFP^ donor, approximately 70% of cells exhibited gene disruption without templated repair (GFP^-^BFP^-^), whereas 25% of cells showed BFP-to-GFP conversion (GFP^+^; Fig. 1B). Repair efficiency was quantified by fluorescence-activated cell sorting (FACS), and the codon-switch validated by sequencing (Fig. 1A, S1A-C). Repair of Cas9-induced breaks using DNA/RNA chimeric donors resulted in lower, albeit still significant, percentages of GFP-positive cells, implicating a reverse transcriptase activity that could synthesize up to 15 rNTPs during DSB repair (Fig.1B). A chimera donor with three scrambled rNTPs did not result in any significant increase in GFP^+^ cells, ruling out random mutagenesis associated with CRISPR/Cas9 editing (Extended Data Fig. 1D). Additionally, RNaseA treatment and gel electrophoresis demonstrated that DNA contamination did not interfere with the assay (Extended Data Fig. 1E).

In a complementary approach, we induced a Cas9 break at the safe harbor genomic *AAVS1* locus and provided donor oligos with a unique three-base pair (bp) insertion (GAT) (Fig. 1C)^28^. The donor was either a pure ssDNA (DNA^1^) or a DNA/RNA chimera donor containing 10 or 22 ribonucleotides spanning the three bp insertion (DNA/RNA^10R^ or DNA/RNA^22R^) (Fig.1D, Supplementary Table 1.2). We assessed repair frequency via next-generation sequencing (NGS) on a 245 bp amplicon flanking the Cas9 cut site. Using CRISPResso^29^, we identified and quantified the RNA insertion as a fraction of the total insertions and deletions (indels). The AAVS1-seq assay detected repair events in the presence of a DNA/RNA chimera, thus corroborating data obtained from the BFP-to-GFP conversion assay (Fig.1D). We further validate these results with droplet digital PCR (ddPCR) using probes that detect insertions at the break site (Fig.S1F-G, Supplementary Table 1.1). We observed a strand bias with DNA/RNA donors compared to ssDNA oligos (Extended Data Fig. 1H-I), similar to previous observations for single-strand templated repair (SSTR) at Cas9-induced breaks^30^, indicating that RNA-containing donors are directly copied at the break site without requiring a double-stranded DNA intermediate. Finally, we examined the repair efficiency of the DNA/RNA^10R^ donor in the presence of a pure DNA donor containing an alternative insertion (CAT, DNA^2^). The competition experiment revealed substantial repair (GAT insertion), though at a reduced efficiency compared to the DNA/RNA^10R^ donor alone (Fig.1D), suggesting competition between the RT-DSBR and SSTR pathways. The two independent assays revealed that human cells possess reverse-transcriptase activity that copies RNA sequences embedded within a single-stranded oligonucleotide to mediate DSB repair.

### RT-DSBR operates independently of LINE-1 retrotransposon and Polθ

To determine whether known human reverse transcriptases participated in RT-DSBR, we investigated the role of LINE-1 retrotransposon and DNA polymerase theta (Polθ). LINE-1 retrotransposons are active in human cells, including HEK293T cells^31,32^, and their mRNA has been detected at sites of DNA damage^33,34^. However, inhibiting LINE-1 reverse transcriptase activity using the HIV reverse transcriptase inhibitors azidothymidine (AZT) or lamivudine (3TC)^35^ did not impair DSB repair as measured by BFP-to-GFP assay and AAVS1-seq, indicating that LINE-1 reverse transcriptase is dispensable for RT-DSBR (Fig. 2A-B, Fig S2A)^36^. We confirmed the efficacy of AZT and 3TC treatment using a fluorescent reporter for LINE-1 reverse transcription activity (Extended Data Fig. 2B)^37^, which showed a three-fold reduction in LINE-1 integration following treatment (Extended Data Fig. 2C).

**Figure 2.**
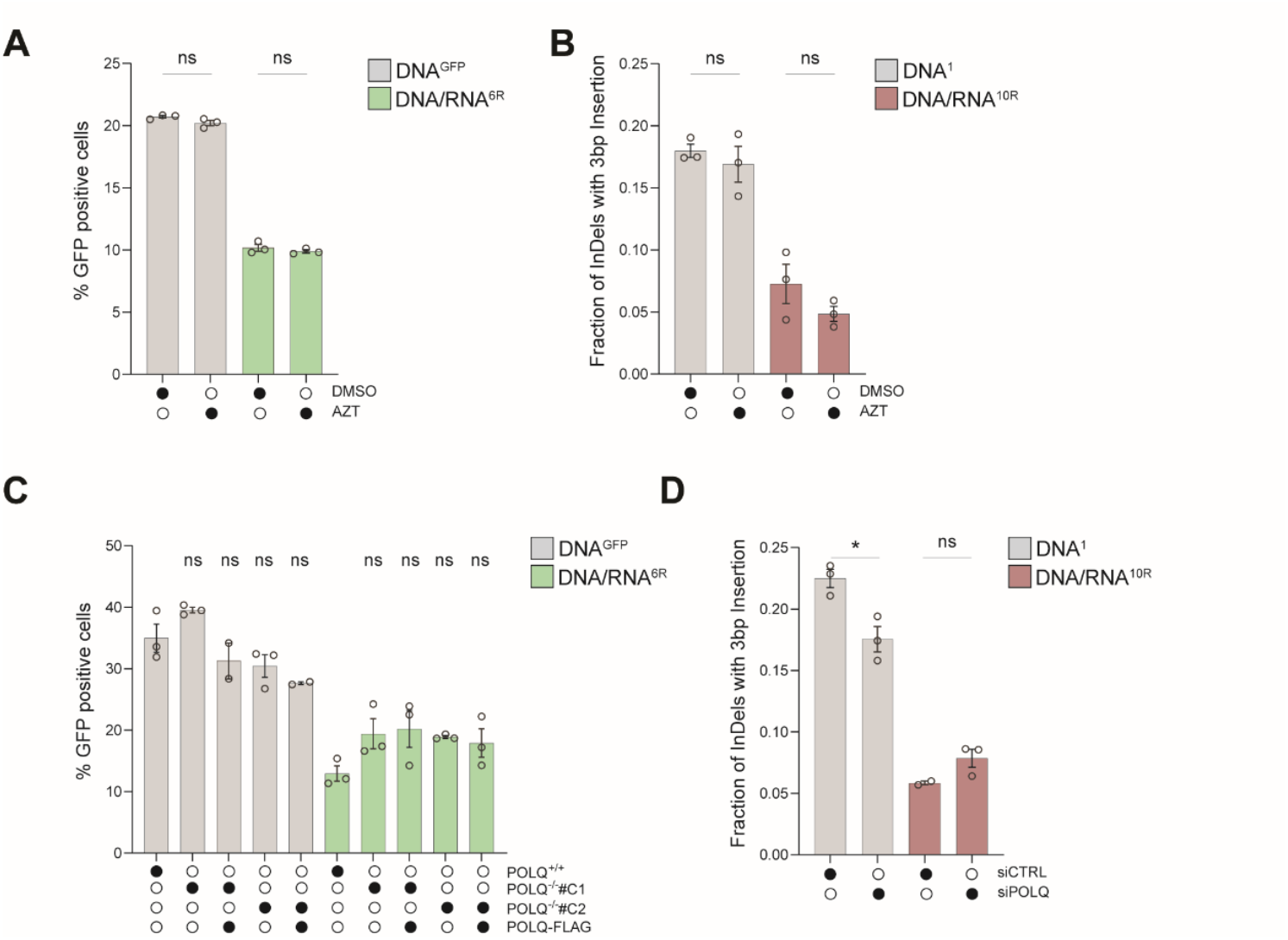
RT-DSBR is independent of LINE-1 and Polθ activity. **A**, BFP-to-GFP assay with DNA^GFP^ and DNA/RNA^6R^ donors in the presence of 10 µM of the HIV reverse transcriptase inhibitor azidothymidine (AZT) or DMSO as a control (n=3). **B**, AAVS1-seq performed with DNA^1^ or DNA/RNA^10R^ donors in the presence of 10 µM AZT or DMSO (n=3). **C**, BFP-to-GFP assay with DNA^GFP^ and DNA/RNA^6R^ donors in two *POLQ*^*-/-*^ clones (#C1 and #C2) with or without complementation by full-length *POLQ* (*POLQ*-FLAG) (n≥2). **D**, AAVS1-seq with DNA^1^ or DNA/RNA^10R^ donors following knockdown of *POLQ* through siRNA, compared to a non-targeting siRNA control (siCTRL) (n≥2). For A-D: Statistical significance was assessed using unpaired Student’s t-test (* p < 0.05). Error bars represent the standard error of the mean (± SEM). See also Figure S2.

A recent study suggested that Polθ, which is critical for MMEJ, has reverse transcriptase activity *in vitro*, and that it can copy a donor template containing two rNTPs *in vivo*^25^. To assess the potential role of Polθ in RT-DSBR, we targeted *POLQ* using CRISPR/Cas9 to generate independent clonally derived *POLQ*^-/-^ cells (Extended Data Fig. 2D-E). BFP-to-GFP assay using DNA^GFP^ and DNA/RNA^6R^ donors revealed that *POLQ*^*-/-*^ cells displayed a similar distribution of repair products compared to *POLQ*^*+/+*^ cells and those rescued with full-length *POLQ*-FLAG (Fig. 2C, S2D-F). In an independent set of experiments, we depleted Polθ using siRNA (Fig. 2D, S2G-H). As expected, Polθ depletion reduced the MMEJ signature following DSB induction^28^ (Extended Data Fig. 2I). Instead, Polθ loss did not impact RT-DSBR, as measured by the BFP-to-GFP reporter and the AAVS1-seq assay (Fig. 2D, S2D-F). Similarly, RT-DSBR was intact in cells treated with the small molecule inhibitor of Polθ, RP6685^38^ (Extended Data Fig. 2J). Based on these findings, we concluded that LINE-1 reverse transcriptase and Polθ activity are dispensable for RT-DSBR.

### A targeted CRISPR/Cas9 screen highlights potential regulators of RT-DSBR

To identify factors that regulate RT-DSBR and to uncover the enzyme responsible for reverse transcription of the RNA moiety, we performed a targeted CRISPR/Cas9 screen using the BFP-to-GFP assay (Fig. 3A). We infected BFP-expressing cells with a focused library of sgRNAs targeting 1,285 DNA damage response (DDR) genes. After ten days, we introduced a Cas9-gRNA RNP complex targeting the BFP locus with either DNA^GFP^ or DNA/RNA^6R^ donors. On day 14, we used FACS to isolate the RT-DSBR-edited GFP^+^ cells from the non-RT-DSBR edited (GFP^−^BFP^−^) cells. We used NGS to determine gRNA abundance in GFP^+^ and GFP^−^BFP^−^populations and applied MAGeCK to identify genes that inhibit RT-DSBR (enriched in GFP^+^) or promote RT-DSBR (depleted in GFP^+^)^39^. By comparing the initial (t=0) and final (t=14) time points, we confirmed stable gene knockdown and a robust hit calling based on the behavior of known essential genes (Extended Data Fig. 3A-B, Supplementary Table 2).

**Figure 3.**
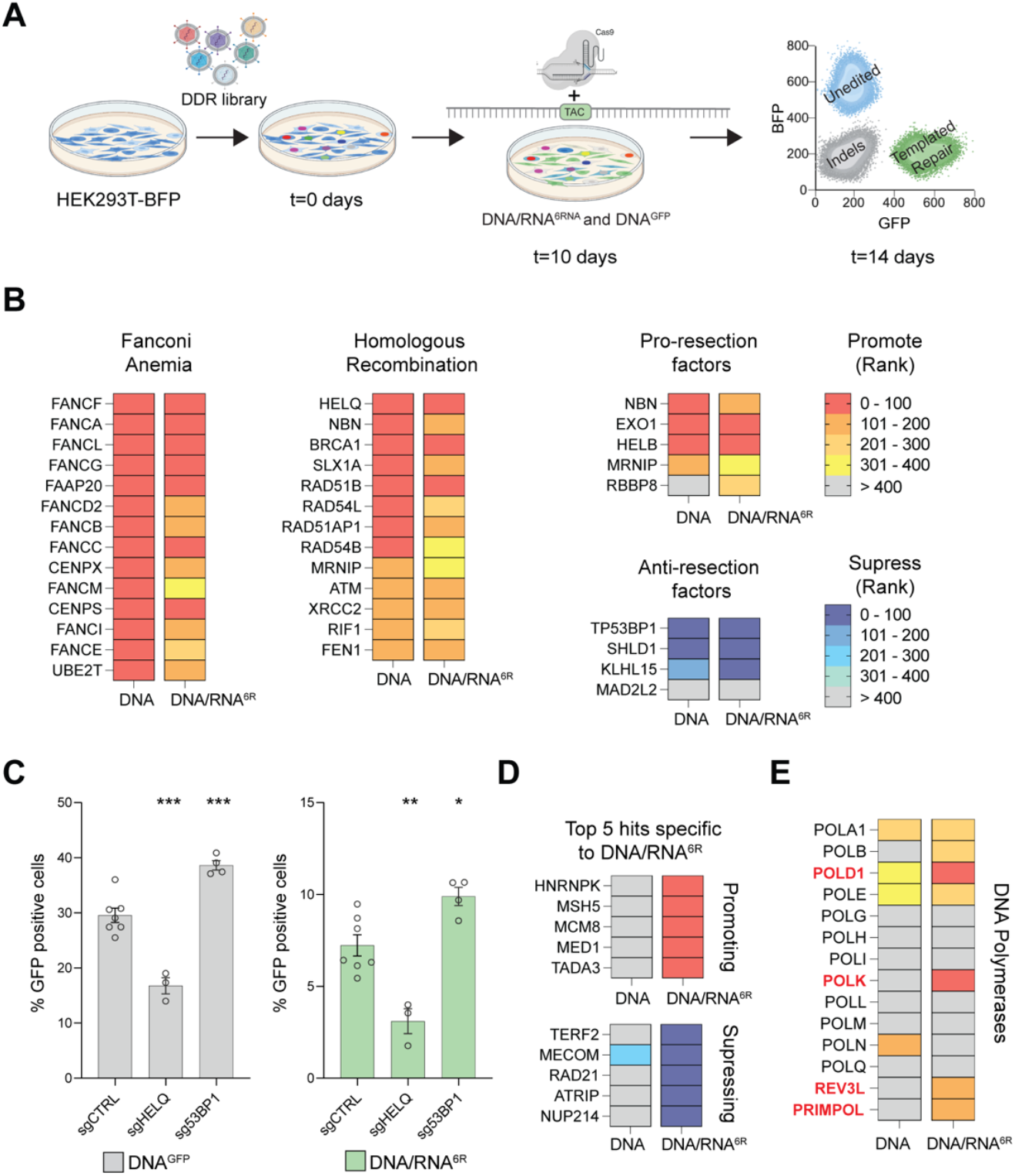
A CRISPR/Cas9 screen identifies factors involved in RT-DSBR. **A**, Schematic representation of a flow-based CRISPR/Cas9 screen performed using the BFP reporter in HEK293T cells. Cells were transduced with Cas9 and sgRNAs from a DNA damage library. After 10 days of sgRNA selection, the BFP-to-GFP assay was carried out using DNA^GFP^ and DNA/RNA^6R^ donors respectively. **B**, The CRISPR/Cas9 screen data were analyzed using the MAGeCK algorithm by comparing the GFP^+^ sorted cells with the GFP^-^ BFP^-^ cells. A heatmap highlights selected genes with high-ranking scores, indicating factors that promote or suppress single-strand template repair. Lower ranks denote stronger hits. **C**, BFP-to-GFP assay results using DNA^GFP^ and DNA/RNA^6R^ donors after knockdown of two top hits that promote (*HELQ*) or suppress (*TP53BP1*) RT-DSBR (n≥3). sgRNA targeting the *AAVS1* locus was used as a control. Statistical significance was assessed using unpaired Student’s t-test (* p < 0.05, ** p < 0.01, *** p < 0.001). Error bars represent the standard error of the mean (± SEM). **D**, Heatmap of the 5 top hits that promote or suppress RT-DSBR. **E**, Comparison of the rank position of major DNA polymerases identified in DNA^GFP^ *vs*. DNA/RNA^6R^ CRISPR/Cas9 screens. See also Figure S3.

We identified several hits that promoted repair using DNA^GFP^ or DNA/RNA^6R^, including factors in the Fanconi anemia pathway (Fig. 3B). We also found that depletion of core factors involved in DNA end-resection (*NBN, EXO1, HELB, BRCA1*) and HR (*HELQ, BRCA1, RAD51B, RAD51AP1*) led to reduced DSB repair with both DNA and DNA/RNA donors (Fig. 3B-C, S3C-D). These findings align with results from a similar CRISPR screen that used a DNA donor^27^, suggesting that both Fanconi anemia and end-resection operate upstream of oligonucleotide-templated repair. Instead, loss of anti-resection factors: *TP53BP1, SHLD1*, and *KLHL15*, resulted in increased templated repair (Fig. 3B-C, S3C-D). The latter observation is consistent with previous studies showing that *TP53BP1* depletion enhanced CRISPR-Cas9 genome editing efficiency^40^. Interestingly, clonally derived *TP53BP1*^*-/-*^ cells showed a three-fold increase in RT-DSBR when using DNA/RNA donors compared to a *TP53BP1*^*+/+*^ cell line and *TP53BP1*^*-/-*^ cells complemented with TP53BP1-FLAG (Fig.S3E-F). In addition to hits that were common to both donor types, we identified genes that uniquely affected DSB repair using the chimera donor (Fig. 3D). Top hits, including hnRNPK and hnRNPC that we validated as RT-DSBR factors using the BFP-to-GFP assay (Fig S3G-H). Given the role of RNA-binding proteins in mRNA maturation and splicing, they may facilitate the retention of the DNA/RNA donor at the site of the break. Alternatively, they may regulate the expression of genes required for RT-DSBR ^41^.

### The translesion polymerase zeta (Polζ) is a reverse transcriptase *in vivo*

To identify the reverse transcriptase responsible for RT-DSBR in human cells, we analyzed DNA polymerases based on their ranking in the CRISPR/Cas9 screen. Specifically, we compared the repair efficiency using the DNA/RNA versus DNA-only donors. Among the 13 human DNA polymerases evaluated, four ranked among the top 200 genes identified in the DNA/RNA donor screen but were less prominent in the DNA-only donor screen (Fig. 3E). These include the catalytic subunit of Polδ (*POLD1*) and Polζ (*REV3L*), *POLK*, and the primase *PRIMPOL*.

To explore the potential reverse transcriptase activity of these polymerases, we performed siRNA-mediated knockdowns and assessed RT-DSBR using the AAVS1-seq assay (Fig. 4A). We target three additional polymerases: Polη (*POLH)*, previously shown to have reverse transcriptase activity *in vitro* and *in vivo*^42–44^, Polμ (*POLM*) which incorporates ribonucleotides at break sites before ligation^45,46^, and Polν (*POLN*) a member of the same family as Polθ^43,47^. Among the seven polymerases tested, knockdowns of *REV3L* and *POLD1* showed a reduction in RT-DSBR (Fig.4A, S4A). However, *POLD1* knockdown also reduced repair events mediated by the DNA donor, indicating that *POLD1* is not specific to RT-DSBR. In contrast, *REV3L* knockdown did not affect repair through the DNA donor, suggesting its specificity in reverse transcribing the RNA template during DSB repair. Consistently, the depletion of *REV3L* led to a significant reduction in RT-DSBR measured by the BFP-to-GFP assay (Fig. 4B, S4B-C). Cell cycle analysis showed no change in the distribution of cells in S-phase following the depletion of *REV3L*, ruling out a cell cycle effect of the knock-down (Extended Data Fig. 4D).

**Figure 4.**
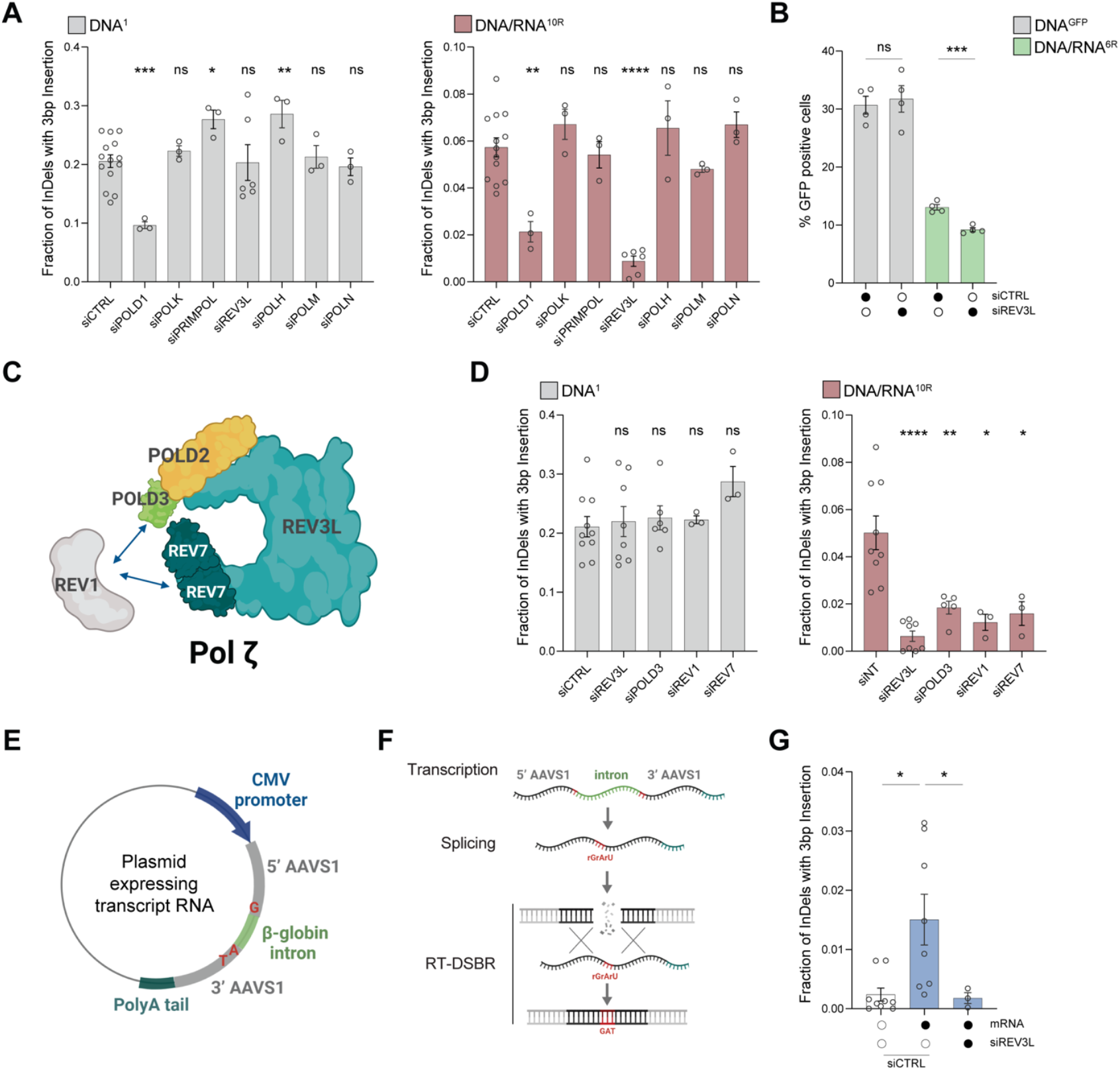
Transcript RNA acts as a donor for polymerase Zeta (ζ) dependent RT-DSBR. **A**, Fraction of repair products from AAVS1-seq using DNA^1^ or DNA/RNA^10R^ donors after siRNA-mediated knockdown of *POLD1, POLK1, PRIMPOL, REV3L, POLH, POLM* or *POLN*, compared to a non-targeting siRNA control (siCTRL). **B**, Percentage of repair products from the BFP-to-GFP assay using DNA^GFP^ or DNA/RNA^6R^ donors following knockdown of *REV3L* with siRNA. **C**, Schematic of the Polζ complex. **D**, Effect of Polζ subunits depletion on AAVS1-seq repair outcomes with DNA^1^ or DNA/RNA^10R^ donors, assessed after siRNA-mediated knockdown. **E & F**, Schematic of a plasmid-based system designated to generate transcript RNA that acts as a donor template. Homology arms (grey) flank the Cas9 break site at the *AAVS1* locus. Light green-β-globin: artificial intron. Darker green: poly-A tail. **G**, Fraction of repair products containing the mutational signature in the presence of no donor or transcript RNA donor, following Cas9-induced breaks. Data were collected after treatment with non-targeting siRNA (siCTRL) (n=7) or siRNA against *REV3L* (n=3). For **A-G**: Statistical significance was assessed using unpaired Student’s t-test, with Welch’s correction in G (* p < 0.05, ** p < 0.01, *** p < 0.001, ****p < 0.0001). Error bars represent the standard error of the mean (± SEM). See also Figure S4 & S5.

Polζ is a multi-subunit complex comprising the catalytic core REV3L and the accessory subunits POLD2, POLD3, and REV7, which interact with the DNA transferase REV1^48^ (Fig. 4C). Consistent with the role of the Polζ complex in reverse transcribing RNA to DNA, depletion of *POLD3, REV1*, and *REV7* subunits led to a reduction in RT-DSBR at the *AAVS1* locus using the DNA/RNA donor (Fig. 4D, S4E). This finding aligns with the CRISPR/Cas9 screen, where the Polζ complex subunits rank higher in the screen using the chimera donor compared to the DNA donor (Extended Data Fig. 4F). Taken together, our results suggest that Polζ is a key reverse transcriptase involved in RT-DSBR.

### Transcript RNA serves as a template for Polymerase ζ-dependent RT-DSBR

Our reporter assays revealed that human cells can utilize synthetic oligonucleotides containing RNA as templates for DSB repair (Fig. 1). This prompted us to investigate whether an RNA transcript also serves as a template for RT-DSBR mediated by Polζ. To that end, we amended the AAVS1-seq assay by introducing an mRNA transcribed from a plasmid as the donor template. This mRNA encodes the *AAVS1* sequence containing a three-base pair (GAT) insertion that is interrupted by the human beta-globin intron^49^ (Fig. 4E-F). The insertion, spanning the splice junction, allowed us to differentiate between repair events via the RNA transcript and those mediated by copying the donor plasmid itself. We verified the correct transcript splicing through PCR analysis using primers spanning the splice site (Extended Data Fig. 4G-H). NGS analysis of the amplicon sequence revealed a small fraction of repair events containing the GAT insertion sequence (Fig. 4G). Significantly, no additional insertion signatures were associated with the presence of the transcript RNA, confirming that the GAT insertion was specific to RT-DSBR activity (Extended Data Fig. 5). When *REV3L* was depleted, the characteristic GAT insertion signature associated with RT-DSBR was significantly reduced (Fig.4G). In conclusion, our data suggest that human cells can use a spliced mRNA complementary to the damage site as a template for DSB repair. Furthermore, this process depends on the Polζ complex, underscoring its essential role in RNA-templated DSB repair.

### Whole intron deletion, a genomic scar reflective of RT-DSBR in human cancers

So far, our experiments have shown that RNA can serve as a template for repairing CRISPR/Cas-9-induced DSBs in human cells. These findings suggest that mRNA transcripts at naturally occurring endogenous break sites might provide a template for DSB repair. However, detecting RT-DSBR at endogenous breaks poses a challenge because RNA-mediated repair typically leaves no detectable scar. An exception would occur if a spliced mRNA transcript were used to repair a break within an intron. In such cases, RT-DSBR could create a distinct signature by precisely removing the intron from the genome, resulting in a whole intron deletion (WID) event (Fig. 5A). Although WIDs are expected to be rare, they are potentially reflective of RT-DSBR activity.

**Figure 5.**
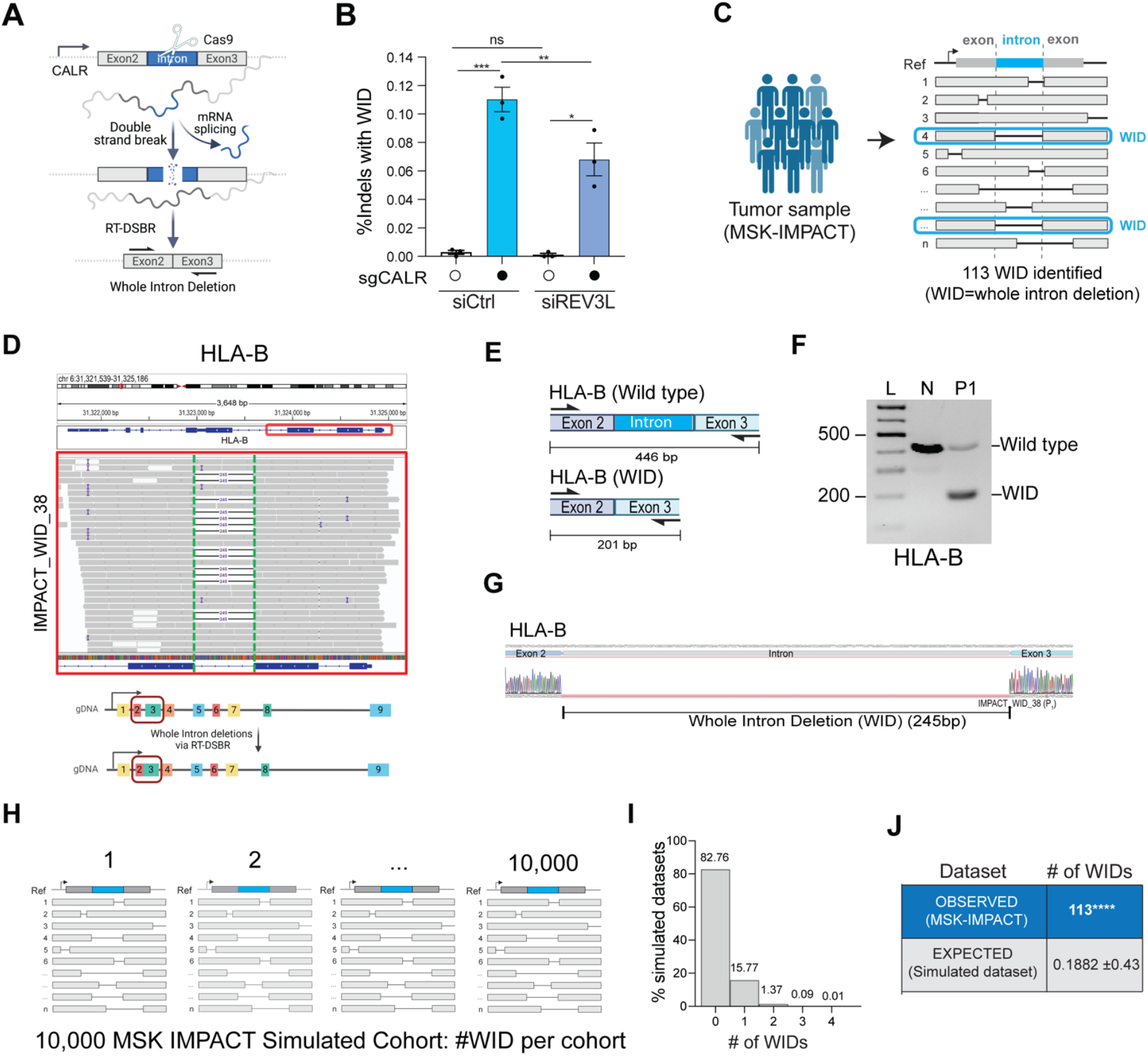
Whole intron deletions from cancer genomes provide *in vivo* evidence of RT-DSBR. **A**, Schematic representation of the CRISPR-Cas9 assay to detect whole intron deletions (WIDs) in human cells. **B**, Quantification of reads containing precise whole intron deletion (WID) of the second intron at the CALR locus in control cells and ones treated with siREV3L (n=3). A CRISPR-Cas9-mediated DSB was introduced at the CALR locus. WIDs driven by endogenous spliced mRNA were measured as a fraction of total repair events. Statistical significance was assessed using an unpaired Student’s t-test (* p < 0.05, ****p < 0.0001). Error bars represent the standard error of the mean (± SEM). **C**, Schematic of the bioinformatic pipeline used to analyze deletions in tumors from the MSK-IMPACT database. The cohort contains 73,030 deletions from 64,544 tumors across all the patient samples. WIDs were identified as deletions that span a precise entire intron. The blue box highlights a read showing perfect intron loss. **D**, Example of a WID found in the *HLA-B* gene of a patient sample from the MSK-IMPACT cohort. Read bases that match the reference are displayed in gray, purple “I” represents insertions, and deletions are indicated with a black dash (–). Alignments displayed with light gray borders and white fill have a mapping quality equal to zero, suggesting they may map to multiple regions across the genome. A 245bp deletion is observed upon targeted NGS that maps precisely to the area corresponding to the intron flanked by Exon 2-3 of the *HLA-B* gene. **E**, Schematic of the exons spanning the WID in *HLA-B* with the flanking primers used to confirm the sequence. **F**, Agarose gel depicting the full-length band corresponding to the locus spanning Exon 2-3 in normal MCF-12A cells (N) and the shorted locus with the intron loss in the tumor sample in HLA-B. P_1_ represents a patient from the MSK-IMPACT cohort. **G**, Sanger sequencing of the PCR products to confirm the presence of the WID in *HLA-B*. **H**, 10,000 MSK-IMPACT-like cohorts were simulated, and the occurrence of WIDs was calculated. Graph representing the number of WID observed in the simulated datasets. **I & J**, Total number of WIDs over 73,030 total deletions identified in 64,544 tumor samples of the MSK-IMPACT database. The number of expected WIDs was calculated after randomization of the deletion locations across the whole genome. Using Fisher’s exact test, empirical P-values were calculated by comparing the observed versus the 10,000 random values (**** p < 0.0001). See also Figure S6.

To investigate whether RT-DSBR can lead to a WID in cells, we targeted a small intron of a highly transcribed gene (CALR)^50^ using a CRISPR-Cas9-induced break in clonally derived *TP53BP1*^*-/-*^ cells. Analysis by TIDE confirmed efficient cleavage at the intron site (Extended Data Fig. 6A). As a positive control, alongside the CRISPR/Cas9 targeting the CALR intron 2, we co-transfected cells with an oligonucleotide designed to mimic a WID event using a donor template. The donor oligonucleotide comprised a DNA/RNA chimera lacking the intronic sequence but complementary to the adjacent exon sequences and containing six ribonucleotides spanning the exon-exon junction. We amplified repair products using primers specific to the flanking exons. Subsequent NGS analysis using CRISPResso2 identified a subset of repaired sequences exhibiting a precise deletion of the second intron (Extended Data Fig 6B – left graph). To test the hypothesis that endogenous spliced CALR mRNA could serve as a repair template, we transfected an sgRNA targeting the CALR intron two without providing an exogenous donor template. We detected a low but statistically significant accumulation of WID events (Fig. 5B and Extended Data Fig. 6B – right graph). Consistent with Polζ promoting RT using mRNA, WID events at the CALR locus were significantly reduced upon depletion of REV3L using siRNA (Fig. 5B and Extended Data Fig. 6C-D). We observed similar WID events driven by endogenous transcripts when targeting another highly transcribed gene – GNAS – with CRISPR/Cas9 at intron 11 (Extended Data Fig. 6E-G), but not the transcriptionally silent gene, IL3, at intron 4 (Extended Data Fig. 6H). These findings suggest RT-DSBR can lead to WID when using spliced mRNA as the template to repair a break.

Next, we examined the repertoire of genomic alterations in cancer genomes from tumor samples, available through MSK-IMPACT and PCAWG^51–53^, to determine whether spliced mRNA could produce intron deletions in tumor cells from naturally occurring endogenous DNA damage. MSK-IMPACT is a hybridization capture-based sequencing assay that analyzes matched tumor/normal samples, covering all coding and selected intronic or regulatory regions of at least 341 essential cancer genes^51^. To identify WIDs, we systematically screened for somatic deletions in 64,544 tumors (from 56,322 patients) that underwent MSK-IMPACT sequencing. By aligning these deletions to the reference genome, we identified 113 unique deletions precisely spanning intronic sequences classified as WIDs (Fig. 5C, Supplementary Table 3.1). As a control, we examined RNA-seq data from the identified tumors, confirming that genes with WIDs are actively transcribed (Extended Data Fig. 6I, Supplementary Table 3.2). We validated the presence of WIDs in two independent genes *(HLA-B and GNAS)* in patient-derived tumor samples through PCR amplification of a region spanning the deleted introns, followed by Sanger sequencing (Fig. 5D-G, S6J-L, Supplementary Table 1.7). To further corroborate our findings from MSK-IMPACT, we conducted an independent analysis using whole-genome sequencing (WGS) data from PCAWG^52^, which contains data from 1,902 patients and tumor samples, with matched normal tissues across 38 tumor types (Extended Data Fig. 6M) ^52^. This analysis revealed 16 additional WIDs, supporting the detection of RT-DSBR activity in a second well-known cancer genome database (Extended Data Fig. 6N-P, Supplementary Table 3.3).

Given the paucity of WID events, it was essential to rule out that their occurrence was due to chance. We conducted a simulation analysis involving 10,000 cohorts of the study genomes, estimating the number of WIDs expected from random deletion events. Each cohort contains a similar number of deletions observed in MSK-IMPACT, with deletions randomly distributed across the genomes while considering deletion lengths and gene content. Although the overall distribution of random deletions closely resembled that pattern seen in MSK-IMPACT, the maximum number of WID events observed across 10,000 simulated cohorts was only four, which occurred in just two cohorts (Fig. 5H-J). These findings suggest that the likelihood of observing 113 WIDs in the MSK-IMPACT data by chance is extremely low (p < 0.0001) (Fig. 5J). We conducted a similar simulation analysis on data from the PCAWG project, which further confirmed that the observed WID events in PCAWG are also unlikely to have occurred by chance (p < 0.0001) (Extended Data Fig. 6O-P).

Furthermore, having observed that of the total 113 WIDs identified in MSK-IMPACT, we found approximately half of the deletions occur in clusters of two or more consecutive WIDs, with some genes losing as many as five consecutive introns (Fig. 6A-B, Table 1, Supplementary Table 3.1). Canonical DSB repair is highly unlikely to lead to the loss of one, let alone sequential introns, as this would require multiple breaks in adjacent introns to occur. The presence of consecutive WIDs provides further evidence that these introns are lost due to using a spliced mRNA as a template, which would lack consecutive introns when used as a donor template. Moreover, our analysis likely underestimates the number of detected genes with >2 consecutive WIDs due to the limitations of the deletion callers in detecting large deletions owing to the sequencing methods used in MSK-IMPACT ^53^. This is observed at genes like XPO1 and JAK1, where we detected two or more consecutive intron losses separated by a single remaining large intron (Table 1). Finally, we confirmed these observations by analyzing adjacent WID events following the CRISPR-mediated cleavage of intron 2 in *CALR*. Notably, we observed a significant accumulation of whole intron deletions events upstream of the break sites (Fig 6C). Our findings implicate RT-DSBR activity in repairing breaks at actively transcribed genes through endogenous spliced mRNA and provide a plausible mechanism for this novel genomic scar (Fig. 6D).

**Figure 6.**
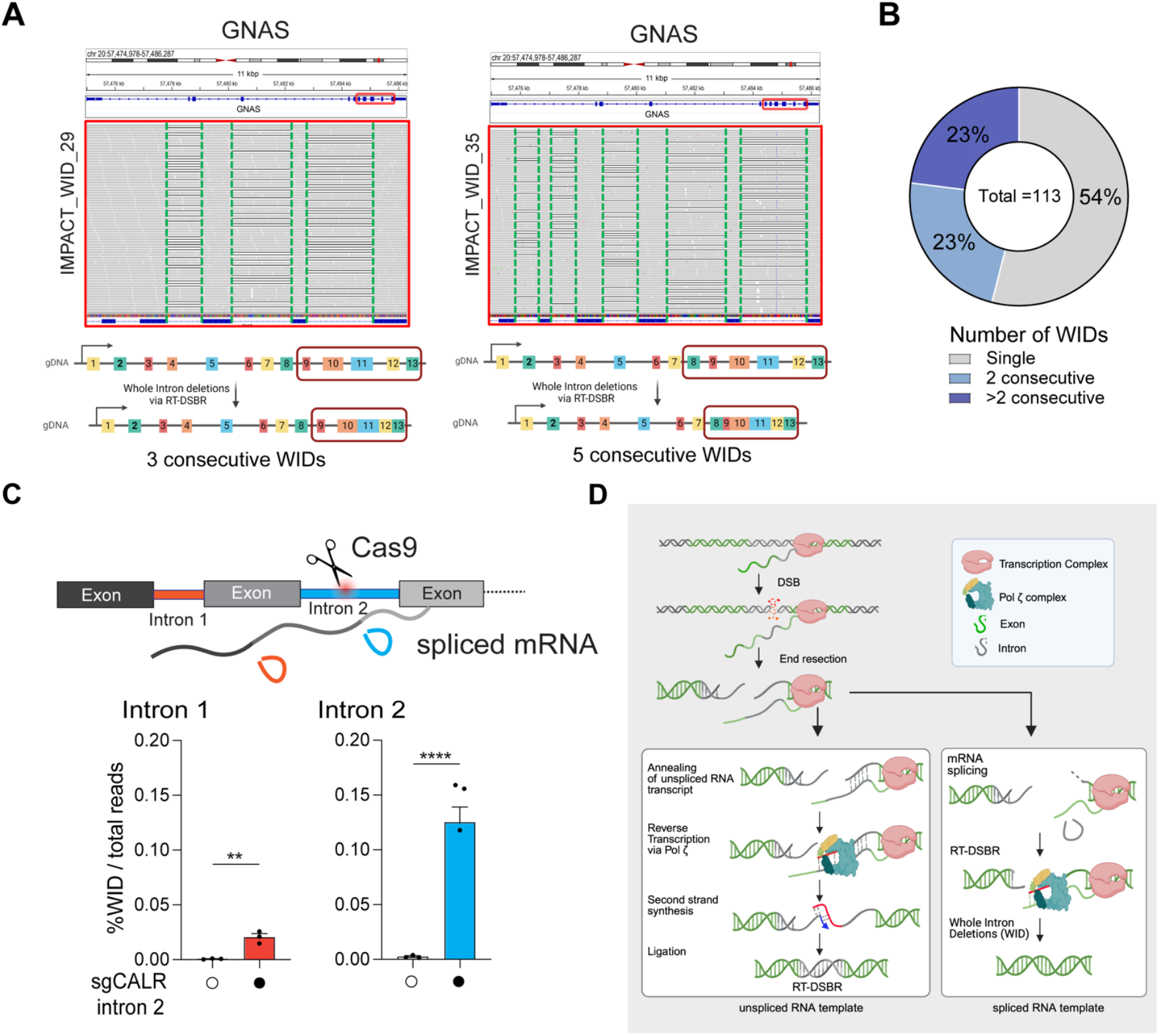
Evidence of Whole Intron deletions in cells. **A**, Example of consecutive WIDs detected in the *GNAS* gene from patient samples sequenced with MSK-IMPACT. Grey bases match the reference genome, and deletions are indicated with a black dash (–). On the left, three deletions were observed at the *GNAS* gene following targeted NGS, precisely mapping the introns flanked by Exon 10-11, 11-12, and 12-13, respectively. On the right, five consecutive deletions were mapped to introns flanked by Exon 8-9, 9-10, 10-11, 11-12, and 12-13. **B**, Frequency of consecutive WIDs observed in the MSK-IMPACT dataset. **C**, Loss of upstream intron following cleavage of CALR intron 2 with CRISPR/Cas9. Top, Schematic representation of multiple introns in CALR gene with cleavage of intron 2. The bottom graph depicts the quantification of reads containing WID in introns adjacent to the cleavage site. Statistical significance was assessed using an unpaired Student’s t-test (* p < 0.05, ****p < 0.0001). Error bars represent the standard error of the mean (± SEM). **D**, Proposed model for RT-DSBR: When a double-strand break (DSB) occurs within an actively transcribed gene, the existing RNA transcript base-pairs with the cleaved template strand and is reverse transcribed by the Polζ complex. The newly synthesized DNA (shown in red) anneals to the resected opposite end, facilitating second-strand synthesis, gap filling, and ligation. The specific polymerase and ligase involved in this process have yet to be identified. If a spliced RNA transcript serves as the repair template, the intronic sequence will be omitted, resulting in a genetic scar known as a whole intron deletion (WID).

**Table 1.** Evidence of consecutive WIDs in tumors. A list of the patient samples exhibiting consecutive WIDs in the same gene, including the locations and sizes of the deletions, is provided.

| Patient ID from this study | Gene | Left exon | Right exon | deletion size observed (bps) | intron size (bps) | Total no of consecutive WIDs |
| --- | --- | --- | --- | --- | --- | --- |
| IMPACT_WID_45 | DICER1 | exon27 | exon26 | 370 | 370 | 2 |
|  |  | exon26 | exon25 | 93 | 93 |  |
| IMPACT_WID_42 | SF3B1 | exon20 | exon19 | 85 | 85 | 2 |
|  |  | exon19 | exon18 | 280 | 280 |  |
|  | SF3B1 | exon17 | exon16 | 216 | 216 | 2 |
|  |  | exon16 | exon15 | 96 | 96 |  |
| IMPACT_WID_26 | XPO1 | exon23 | exon22 | 417 | 417 | 2 |
|  |  | exon22 | exon21 | 845 | 845 |  |
| IMPACT_WID_36 | PIK3CA | exon18 | exon19 | 561 | 561 | 2 |
|  |  | exon19 | exon20 | 103 | 103 |  |
| IMPACT_WID_32 | JAK1 | exon25 | exon24 | 738 | 738 | 2 |
|  |  | exon24 | exon23 | 591 | 591 |  |
|  | JAK1 | exon22 | exon21 | 360 | 360 | 2 |
|  |  | exon21 | exon20 | 1013 | 1013 |  |
| IMPACT_WID_05 | ERRFI1 | exon4 | exon3 | 911 | 911 | 2 |
|  |  | exon3 | exon2 | 110 | 110 |  |
| IMPACT_WID_11 | CSDE1 | exon13 | exon12 | 877 | 877 | 2 |
|  |  | exon12 | exon11 | 596 | 596 |  |
| IMPACT_WID_21 | DAXX | exon7 | exon6 | 194 | 194 | 2 |
|  |  | exon6 | exon5 | 160 | 160 |  |
| IMPACT_WID_20 | CSDE1 | exon13 | exon12 | 877 | 877 | 2 |
|  |  | exon12 | exon11 | 596 | 596 |  |
| IMPACT_WID_12 | GNAS | exon11 | exon12 | 252 | 252 | 2 |
|  |  | exon12 | exon13 | 281 | 281 |  |
| IMPACT_WID_04 | CDK8 | exon10 | exon11 | 718 | 718 | 2 |
|  |  | exon11 | exon12 | 118 | 118 |  |
| IMPACT_WID_34 | GNAS | exon10 | exon11 | 146 | 146 | 3 |
|  |  | exon11 | exon12 | 252 | 252 |  |
|  |  | exon12 | exon13 | 281 | 281 |  |
| IMPACT_WID_29 | GNAS | exon10 | exon11 | 146 | 146 | 3 |
|  |  | exon11 | exon12 | 252 | 252 |  |
|  |  | exon12 | exon13 | 281 | 281 |  |
| IMPACT_WID_22 | GNAS | exon10 | exon11 | 146 | 146 | 3 |
|  |  | exon11 | exon12 | 252 | 252 |  |
|  |  | exon12 | exon13 | 281 | 281 |  |
| IMPACT_WID_09 | GNAS | exon10 | exon11 | 146 | 146 | 3 |
|  |  | exon11 | exon12 | 252 | 252 |  |
|  |  | exon12 | exon13 | 281 | 281 |  |
| IMPACT_WID_26 | XPO1 | exon16 | exon15 | 126 | 126 | 4 |
|  |  | exon15 | exon14 | 85 | 85 |  |
|  |  | exon14 | exon13 | 166 | 166 |  |
|  |  | exon13 | exon12 | 840 | 840 |  |
| IMPACT_WID_19 | GNAS | exon8 | exon9 | 97 | 97 | 5 |
|  |  | exon9 | exon10 | 104 | 104 |  |
|  |  | exon10 | exon11 | 146 | 146 |  |
|  |  | exon11 | exon12 | 252 | 252 |  |
|  |  | exon12 | exon13 | 281 | 281 |  |
| IMPACT_WID_35 | GNAS | exon8 | exon9 | 97 | 97 | 5 |
|  |  | exon9 | exon10 | 104 | 104 |  |
|  |  | exon10 | exon11 | 146 | 146 |  |
|  |  | exon11 | exon12 | 252 | 252 |  |
|  |  | exon12 | exon13 | 281 | 281 |  |

## DISCUSSION

Emerging evidence suggests that RNA transcripts can indirectly shape the landscape of DSB repair by modulating three canonical repair pathways: HR, NHEJ, and MMEJ^17,54^. Transcription and DNA repair are intrinsically linked processes, as evident by the evolution of transcription-coupled nucleotide excision repair, a specialized DNA repair pathway^55^. Moreover, the annealing of RNA with the complementary strand of DNA to form an R-loop can act as a scaffold that recruits repair factors and increases HR efficiency. This effect is pronounced in highly transcribed genes, thus providing evidence of a role for RNA in modulating the outcome of DSB repair^16,55^. Despite this, understanding whether RNA can directly serve as a template for DSB repair has been challenging due to the lack of tools to assess its contribution in higher eukaryotes. In this study, we demonstrate that RNA serves as a template for DSB repair via reverse transcription facilitated by the DNA polymerase ζ complex. We show that RT-DSBR using mRNA is a rare mutagenic pathway in human tumors, with a highly characteristic WID genomic scar. Given the abundance of RNA and its encoding of genetic information, utilizing RNA to restore lost genetic information following DSBs may be potentially driven by selective pressure to preserve the integrity of highly transcribed genes.

### Transcript RNA as a template for DSB repair in human cells

Based on our findings, we propose a model in which DSBs occurring in actively transcribed genes can utilize the corresponding RNA transcript as a template for repair (Fig. 6D). It remains unclear whether the RNA transcript used for repair is generated before or after DSB formation. However, since transcription is disrupted in response to DNA damage^4–6^, we favor a scenario in which the donor RNA template is transcribed before DSB formation. Once the RNA anneals to the processed DNA end, we demonstrate that Pol ζ can use the mRNA to fill the gap via reverse transcription, restoring the original genetic information.

Recently, other translesion polymerases, specifically Polη and Polθ, have exhibited reverse transcriptase activity in both *in vitro* and *in vivo* settings^25,44^. However, our reporter assays could not detect reverse transcription activity for Polη and Polθ (Figs. 2&4). In addition to Pol ζ, our CRISPR/Cas9 screen identified 53BP1 as a factor that suppresses RT-DSBR, which we validated in a 53BP1^-/-^ clone. Given that 53BP1 is known to counteract DNA resection^56^, this implies that resection may be crucial step required to process the breaks before the RNA template can anneal. This genetic manipulation, which increases the use of RT-DSBR, will be an important tool for the dissection of other factors involved in this pathway.

### Conservation of RNA-templated DSB repair from yeast to humans

RT-DSBR appears to be a conserved mechanism from yeast to humans^18,19^. In *S. cerevisiae*, Ty1 retrotransposons mediate cDNA synthesis from mRNA for DSB repair *via* an HR-like mechanism. In the absence of Ty1, Polζ reverse transcribes the RNA at break sites in cis to mediate repair^20^. Unlike yeast, LINE-1 reverse transcriptase is dispensable for RT-DSBR in human cells (Fig. 2). Instead, Polζ has a prominent role in copying the RNA to mediate DSB repair. Furthermore, as opposed to RT-DSBR in yeast, which was detected only in the absence of RNaseH1 and RNaseH2, we detect low but significant RNA templated repair in human cells competent for both enzymes (Fig. 1, Fig. 4D, G). While these results suggest that RNA:DNA hybrid removal is not essential for RT-DSBR in human cells, whether RNaseH1 and RNaseH2 have a role in this process remains unexplored.

Polζ is a critical translesion synthesis (TLS) polymerase responsible for synthesizing across various types of DNA lesions, including abasic sites and UV-damaged bases^48^. In contrast to other TLS polymerases, Polζ belongs to the B-family of DNA polymerases, which includes accurate replicative polymerases. However, Polζ lacks 3′-5′ exonucleolytic proofreading activity, contributing to spontaneous mutagenesis in eukaryotic cells^57^. Notably, Polζ was reported to bypass single ribonucleotides in yeast, preventing replication fork stalling. *In vitro* studies have shown that the catalytic subunit of Polζ can efficiently bypass four ribonucleotides in tandem, highlighting its potential reverse transcriptase activity^58,59^. Deleting *REV3L* in chicken or mammalian cells causes hypersensitivity to genotoxic stress, including agents that induce DSBs^60,61^. Our findings highlight a novel function of Polζ to copy RNA into DNA during RT-DSBR. The mechanisms by which Pol ζ is recruited to DSBs and regulated at these sites remain unknown. Additionally, future efforts exploring whether transcription influences its recruitment to DSBs may provide further insights into its role in RT-DSBR.

Interestingly, in yeast, the Ty1 retrotransposon mediates the synthesis of complementary DNA from mRNA, which can be subsequently used for DSB repair through a homologous recombination-like mechanism^19^. However, according to our reporter assays, the main active human transposons, LINE1, do not play a role in this process (Fig. 2). In contrast, in both yeast and human cells, Pol ζ appears to directly reverse transcribe mRNA at the site of the DSB (Fig. 4).

### Whole Intron Deletion: a genomic signature of RT-DSBR

Our model predicts that DSB repair can occur without leaving any detectable scar when pre-spliced RNA transcripts are used as templates. As such, detecting RT-DSBR activity in higher eukaryotes is particularly challenging because, in most cases, it leaves no genomic signature. However, when the RNA donor has already undergone splicing, repair of a break within an intron would prompt its elimination from the genome. In such cases, reverse transcription of the spliced RNA could result in genetic scars, such as WID, providing evidence of RT-DSBR activity in vivo. We provide evidence of intron loss by inducing a Cas9-break in highly transcribed genes (Fig. 5A-B). Importantly, we provide the first evidence of the accumulation of WID in human tumor samples, indicating that DSBs can be repaired using spliced mRNA. The low frequency of WIDs in tumors limits our ability to determine whether specific mutations or genomic features influence this pathway and contribute to intron loss. The detection of a cluster of 2 or more consecutive and precise WIDs (Fig. 6) strongly indicates the use of RT-DSBR. This scenario can only be explained by spliced mRNA serving as a template for RT-DSBR, especially since other repair mechanisms are highly unlikely to result in the loss of sequential introns with precise exon-exon junctions.

Although WIDs are rare in tumors, we cannot exclude the possibility that intron loss events also occur in normal cells. Phylogenetic studies comparing genomes of organisms with abundant introns to those with fewer introns reveal a bias towards 3’ end intron loss. Two primary hypotheses have been suggested to explain this bias: one theory, based on studies in *C. elegans* and *D. melanogaster*, posits that intron loss results from error-prone DSB repair by MMEJ and is driven by sequence homology near the ends of the break site^62^. An alternative hypothesis suggests that intron loss is due to retrotransposon-mediated reverse transcription of spliced mRNA^63–65^. Our data suggest that neither human retrotransposon activity nor MMEJ is involved in RT-DSBR-dependent intron loss. Instead, we show that Polζ-mediated RT-DSBR is active in human cells and can produce intron loss, both spontaneously and in response to a break in an intron. Whether RT-DSBR contributed to intron loss during evolution remains to be determined. In a related context, it has been suggested that some pseudogenes form when mRNA transcripts are reverse-transcribed by LINE-1 and integrated into new locations in the genome^66^. These processed pseudogenes lack introns and may be driven by RT-DSBR activity in the germline.

### RNA-templated DSB repair in physiological conditions

With ~78% of the human genome actively transcribed^3^, RT-DSBR may be more common in transcriptionally active loci, especially in contexts where homologous DNA templates are absent. Specifically, RT-DSBR may offer an error-free repair system for active genes in non-dividing cells, where error-free HR is blocked, and NHEJ becomes the only available option for DSB repair. For example, in neuronal cells, topoisomerase II-induced DSBs are stimulated by neuronal activity to resolve topological constraints at highly transcribed genes^67,68^. In such settings, RT-DSBR may offer a safer alternative to NHEJ for repairing these physiological breaks in non-dividing cells, thus safeguarding the genome. Selective pressures may have favored the development of such mechanisms to maintain genomic integrity, for example, at highly expressed loci, which are vital for cellular homeostasis.

In summary, our findings provide new insights into the role of transcript RNA as a template for DSB repair, highlighting a novel connection between transcription and DSB repair in human cells. Further studies in diverse biological contexts will unravel the full spectrum of RT-DSBR activity and its implications for genome stability and evolution.

## AUTHOR CONTRIBUTION

M.J., A.B., S.N.P., and A.S. conceived the experimental design and implemented the study. A.B. performed the BFP-to-GFP assay with help from H.S. (Fig. 3 & Fig. S3) and J.W. (Fig. S1a), and M.J. performed the AAVS1-seq assay with help from N.M.D. and J.G-A. (Fig. 4) and K.S.A. (Fig 2). A.D. performed the CRISPR screen together with A.B., and H.S. helped with the screen analysis. S.A.-S., S.H., and D.H. helped with the AAVS1-seq assay optimization. M.J., N.M.D., J.P., Y.Z., A.G., T.Y., P.C.B., N.R. and J.S.R.-F. Contributed to the computational analysis; J.P., Y.Z. and N.R. analyzed the CRISPResso data; J.P. developed the computational pipeline to search for tumor-specific WIDs with help from Y.Z., T.N.Y., P.S., and A.N-G., while A.G. developed the mathematical modeling. H.S. optimized and performed the intron-loss assay in cells. B.A. in E.L-D. lab generated the DDR-focused CRISPR/Cas9 library. The manuscript was prepared by M.J. and A.B. and revised by N.M.D., H.S., S.N.P., and A.S. with input from all authors.

## ACKNOWLEDGEMENT

We acknowledge using the Integrated Genomics Operation Core, funded by the NCI Cancer Center Support Grant (CCSG, P30 CA08748), Cycle for Survival, and the Marie-Josée and Henry R. Kravis Center for Molecular Oncology. Illustrations were created with BioRender.com. We thank Erik Anderson for helping with designing the RNA-donor plasmid. We thank Ronglai Shen for the helpful discussions on developing the simulations. We thank Martin Stojaspal for technical support with the native PAGE. We thank Raj Chari and Genome Modification Core of NCI for assistance with synthesizing the DDR-focused CRISPR/Cas9 library. We acknowledge the Powell and Sfeir lab members for commenting on the manuscript. This work is supported by grants from NIH/NCI (R01CA229161 and U01CA231019) for A.S. M.J. is supported by the AACR-Swim Across America Cancer Research Fellowship (20-40-64-JALA). S.N.P. is funded by the Breast Cancer Research Foundation and NIH/NCI P50 CA247749. J.S.R.-F. is funded in part by the Breast Cancer Research Foundation, by a Susan G Komen Leadership grant, and by the NIH/NCI P50 CA247749 grant. P.C.B. is supported by the NCI awards P30CA016042, R01CA244729, and U2CCA271894.

## EXTENDED DATA FIGURES

**Extended Data Figure 1.**
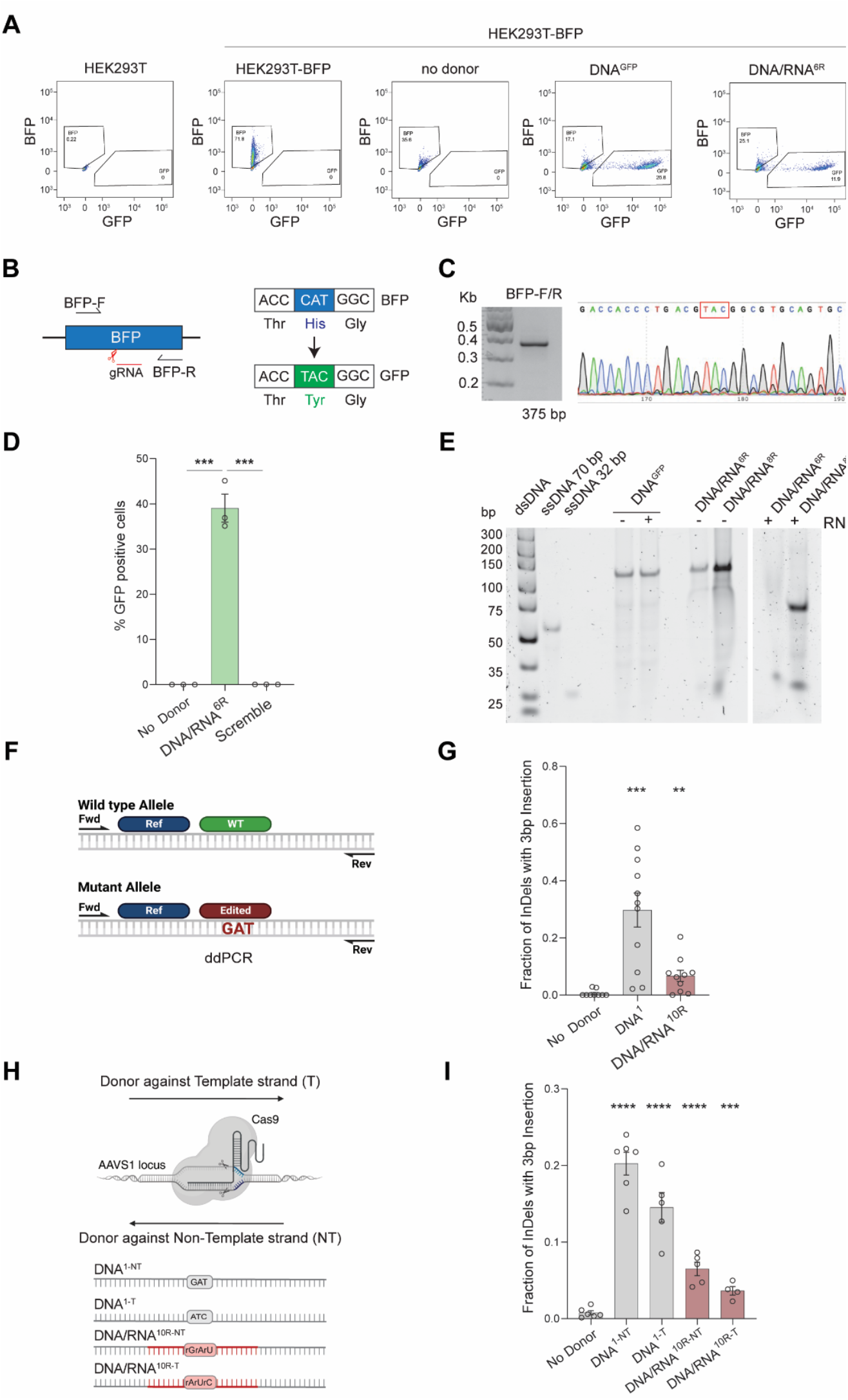
Human cells use RNA-containing oligos to repair DSBs. **A**, Examples of FACS gating for BFP-to-GFP assay. **B-C**, Schematic of the BFP-to-GFP assay showing the CAT to TAC mutation that induces the BFP to GFP switch. Arrows indicate the primers used to amplify the region around the swap codon, and the resulting PCR band of 375 bp is shown on the left. Sanger sequencing profile confirms the mutated population. **D**, BFP-to-GFP assay using a DNA/RNA^6R^ donor containing a scramble sequence in the RNA portion instead of the amino acid switch codon in clonally derived *TP53BP1*^*-/-*^ cells described in Extended Data Fig. 3E-F (n=3). **E**, Native PAGE gel showing the purity of donors used in Fig.1B. Double-stranded (dsDNA) and single-stranded (ssDNA) were loaded as reference. The DNA/RNA^6R/8R^ donors were cleaved with RNaseA and the DNA^GFP^ donor was used as a negative control. **F**, Schematic of the probes and primers used for the ddPCR to detect the wild type (WT) or mutant allele (Edited) in the AAVS1-seq assay. **G**, Quantification of the relative fraction of droplets with 3bp insertion measured with ddPCR after AAVS1-seq (n=10). **H**, Schematic of AAVS1-seq assay using as donors either the template strand (DNA^1-T^ and DNA/RNA^10R-T^) or the non-template strand (DNA^1-NT^ and DNA/RNA^10R-NT^). **I**, AAVS1-seq, measures the percentage of repair products containing the mutational signature after the Cas9 DSB is repaired in the presence of no donor or the donors from panel G (n≥4). For **D, G, I**: Statistical significance was assessed using unpaired Student’s t-test (* p < 0.05, ** p < 0.01, *** p < 0.001, ****p < 0.0001). Error bars represent the standard error of the mean (± SEM).

**Extended Data Figure 2.**
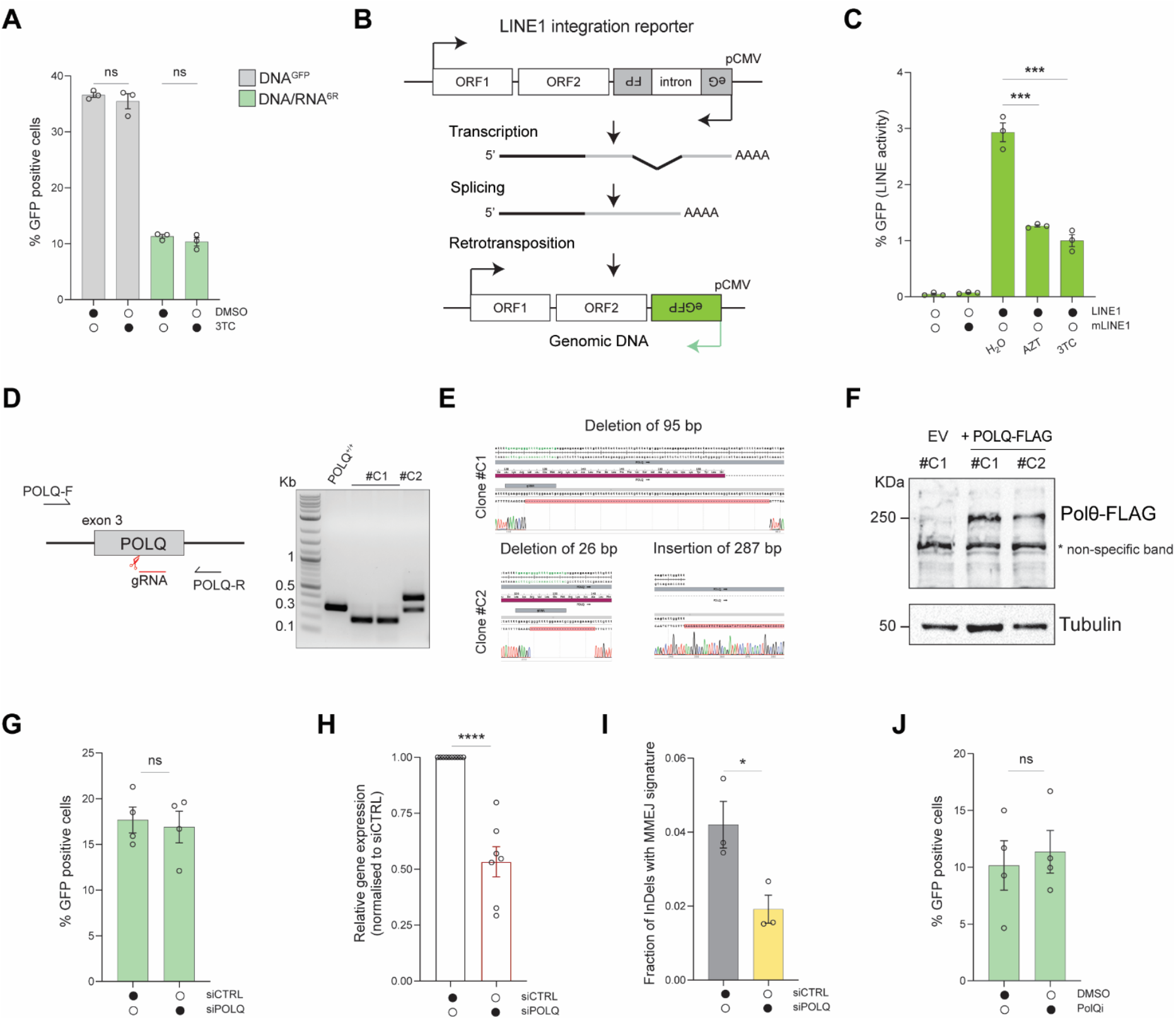
RT-DSBR is independent of LINE-1 and Polθ activity. **A**, BFP-to-GFP assay with DNA^GFP^ and DNA/RNA^6R^ donors in the presence of 10 µM of the HIV reverse transcriptase inhibitor lamivudine (3TC) or DMSO as a control (n=3). **B**, A plasmid encodes a full-length, retrotransposition-competent LINE-1 element driven by its natural promoter. An EGFP retrotransposition reporter cassette is inserted in the LINE-1 3′ UTR, with the EGFP gene in reverse orientation to the LINE-1 sequence. The sequence is interrupted by an intron in the same orientation as LINE-1. Transcription from the LINE-1 promoter produces an mRNA that does not express EGFP due to the reverse orientation, but if retrotransposition occurs and LINE-1 integrates into the genome, a correctly oriented EGFP mRNA is transcribed from the CMV promoter, resulting in EGFP expression. A control plasmid (mLINE-1), containing a point mutation in ORF1 to disable retrotransposition, is used as a negative control. **C**, HEK293T-BFP cells were incubated with 10 µM of AZT or 3TC 24 hours before transfection with the LINE-1 retrotransposition reporter plasmid and kept in cells throughout the experiment. Transfectants were selected with puromycin and analyzed by flow cytometry five days post-transfection (n=3). **D**, Schematic of *POLQ* knock-out strategy. DNA gel for genotyping PCR showing the results of CRISPR-Cas9 targeting in two clones. *POLQ*^*+/+*^ untargeted cells were used as a negative control. **E**, Clones #C1 and #C2 showed insertion and deletion by Sanger sequencing. **F**, Western blot analysis of POLQ^-/-^ clones and HEK293T-BFP cells transfected with full-length POLQ-FLAG plasmid. Tubulin was used as a loading control. **G**, BFP-to-GFP assay with DNA/RNA^6R^ donor after knockdown of *POLQ* with siRNA (n=4). **H**, qPCR analysis of *POLQ* mRNA expression after siRNA knockdown of the samples analyzed in main Fig. 2D and Extended Data Fig. 2G (n=7). Relative gene expression was normalized using *ACT1* as a housekeeping gene. **I**, Relative fraction of indels with microhomology-mediated end-joining signature (12 bp deletion with five bp microhomology) after knockdown of *POLQ* through siRNA in the AAVS1-seq assay performed in main Fig. 2D (n=3). **J**, BFP-to-GFP assay with DNA/RNA^6R^ donor after treatment with the Polθ inhibitor RP6685 (n=3). For **A, C, G-J**: Statistical significance was assessed using unpaired Student’s t-test (* p < 0.05, ** p < 0.01, *** p < 0.001, ****p < 0.0001). Error bars represent the standard error of the mean (± SEM).

**Extended Data Figure 3.**
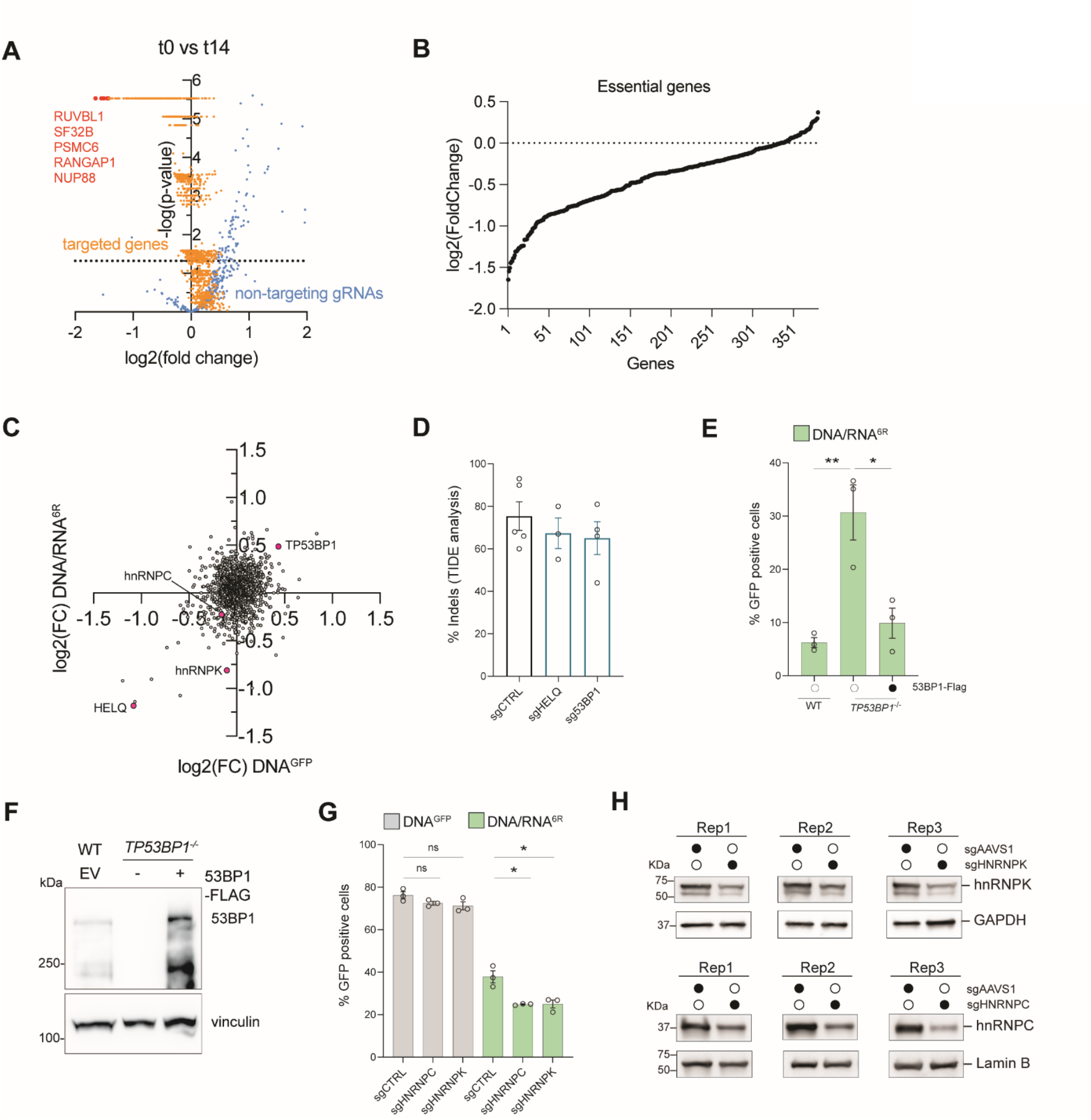
A CRISPR/Cas9 screen identifies factors involved in RT-DSBR. **A**, Volcano plot depicting genes from the DNA repair library enriched or depleted in the CRISPR/Cas9 screen and gRNA controls. Representative essential genes are highlighted in red, negative control untargeted gRNAs are shown in blue, and targeted genes are in orange. Data were generated from two independent BFP-to-GFP assays performed on the same infected cells: n=2 technical duplicates. **B**, Plot representing log2 fold change in the CRISPR/Cas9 screen comparing t=0 versus t=14 days for 360 genes in the DDR library considered as “common essential”^69^. **C**, Comparison of the log2 fold change for each gene in the CRISPR/Cas9 screen with the DNA/RNA^6R^ or the DNA^GFP^ donors. **D**, Editing efficiency calculated with TIDE analysis^70^ for sgRNAs in Figure 3C (n≥3). **E**, BFP-to-GFP assay in a *TP53BP1*^*-/-*^ clone rescued by 53BP1 overexpression (53BP1-FLAG) (n=3). **F**, Western blot of cells deleted for *TP53BP1* (second lane) and complemented with 53BP1-FLAG (third lane). **G**, BFP-to-GFP assay in cells depleted for hnRNPK or hnRNPC in *TP53BP1*^*-/-*^ cells via the DNA^GFP^ and the DNA/RNA^6R^ donors (n=3). **H**, Western blot of cells depleted for *HNRNPK* and *HNRNPC* with sgRNAs (n=3). For **D, E, G**: Statistical significance was assessed using unpaired Student’s t-test (* p < 0.05, ** p < 0.01). Error bars represent the standard error of the mean (± SEM).

**Extended Data Figure 4.**
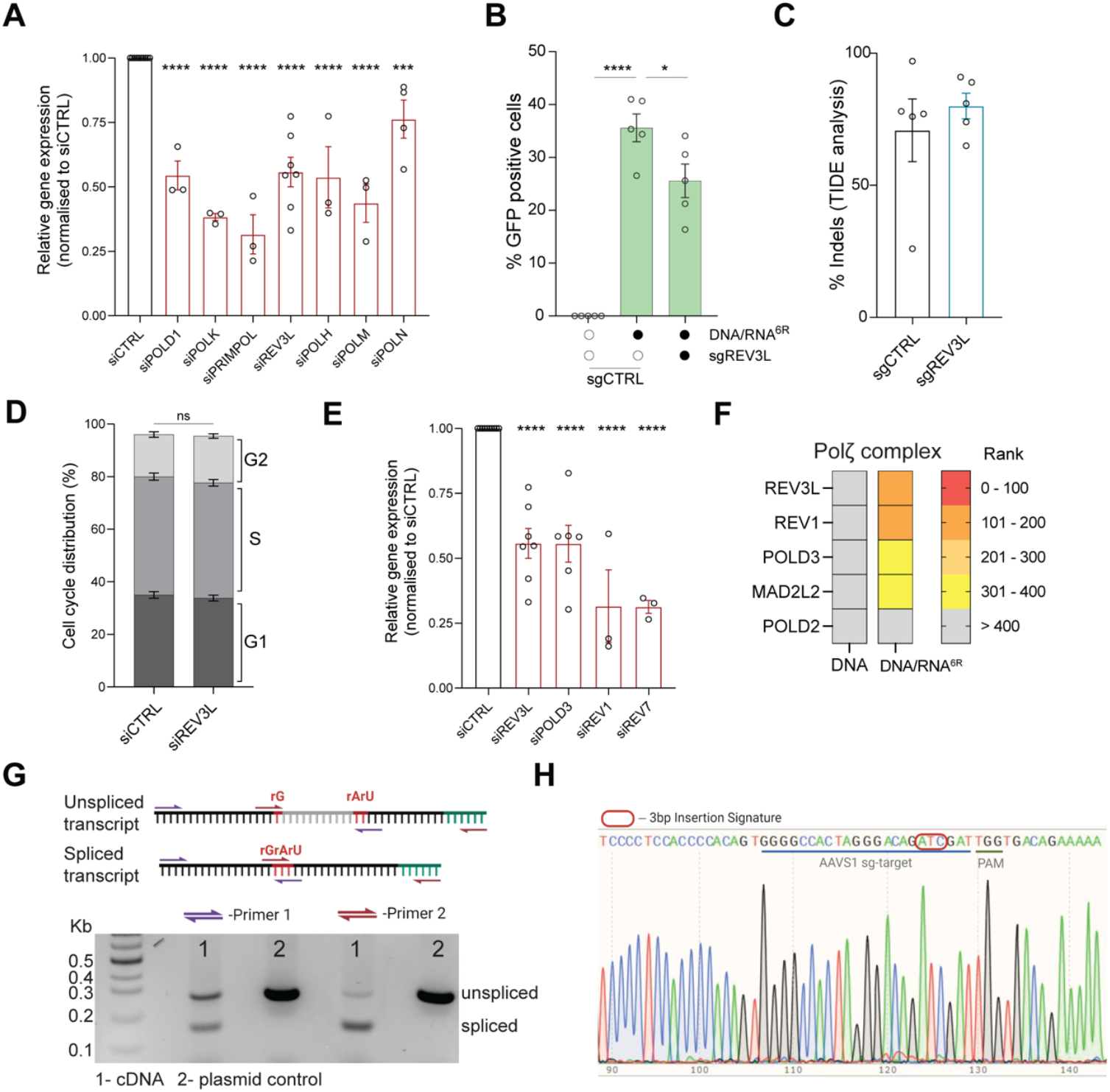
The TLS DNA polymerase zeta (ζ) is a reverse-transcriptase *in vivo*. **A**, Gene expression analysis by qPCR for siRNA knockdown of the major DNA polymerases in Fig. 4A. *ACT1* was used as a housekeeping gene (n≥3). **B**, Effect of *REV3L* targeting via sgRNA on the BFP-to-GFP assay using the DNA/RNA^6R^ donor in the *TP53BP1*^*-/-*^ clone (n=5), compared to a sgRNA control (sgAAVS1). **C**, Editing efficiency of sgRNA against *REV3L* in Extended Data Fig. 4B calculated with TIDE (n=5). **D**, Comparison of the cell cycle distribution of cells treated with and without si*REV3L* (n=5). **E**, Gene expression analysis by qPCR of siRNA knockdown of the subunits of Polζ analyzed in Fig. 4D. *ACT1* was used as a housekeeping gene. Student’s t-test was run for statistical significance. Error bars represent the standard error of the mean. **F**, Heatmap of Polζ subunits showing their rank in the screen. **G**, Schematic of the transcribed RNA with the location of the primer pairs used to validate splicing of the transcript donor RNA containing the 3bp mutation signature (See Methods). Gel image showing the spliced and unspliced RNA species amplified from (1) cDNA from cells expressing the plasmid that codes for the donor RNA. (2) Plasmid control with band corresponding to the unspliced RNA product. **H**, Sanger sequencing profile of the spliced species containing the mutational signature is shown. For A, B, D and E, Statistical significance was assessed using unpaired Student’s t-test (* p < 0.05, ** p < 0.01, *** p < 0.001, **** p < 0.0001). For D: Statistical significance was assessed using 2-way ANOVA. Error bars represent the standard error of the mean (± SEM).

**Extended Data Figure 5.**
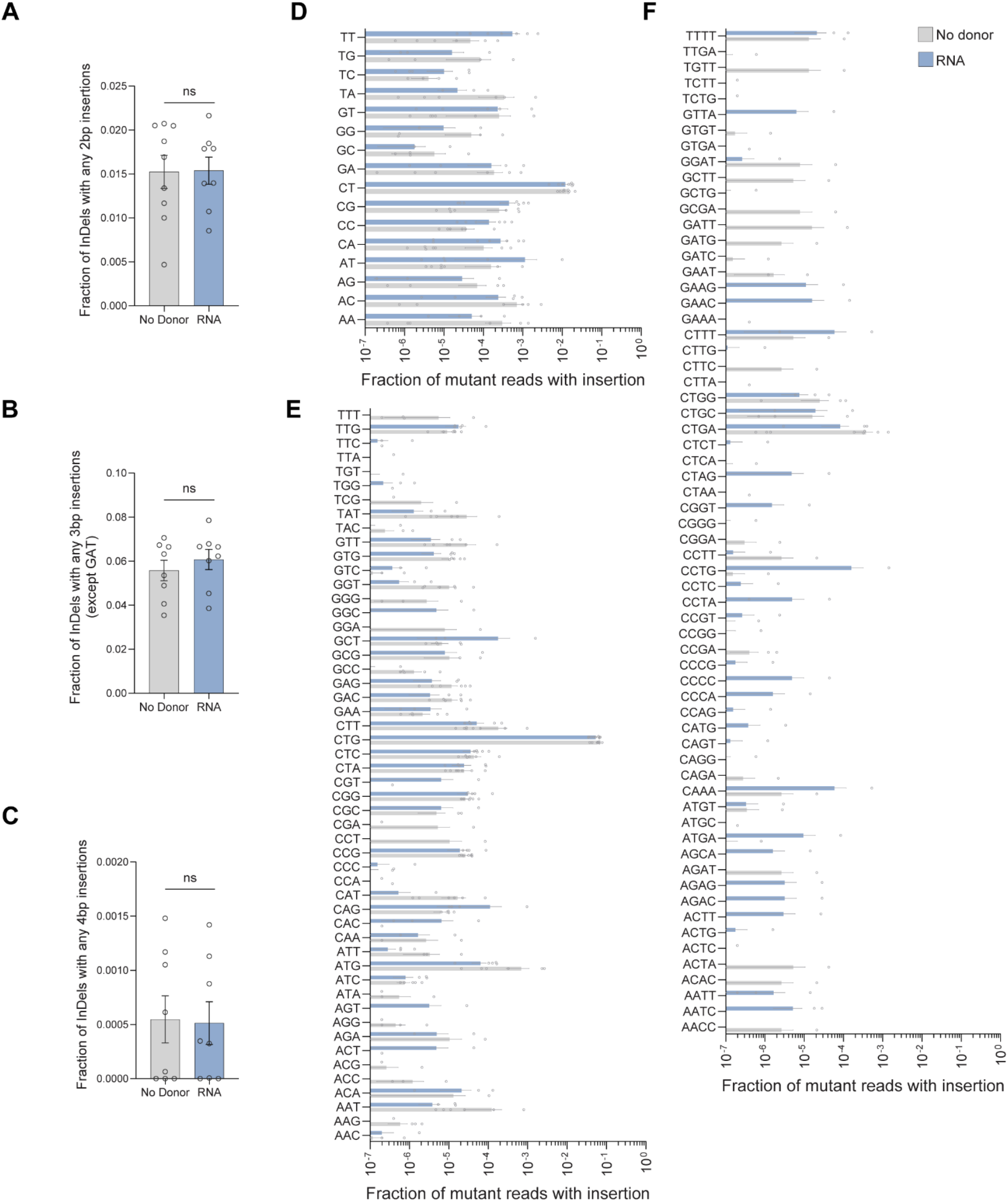
Repair via transcript RNA as a donor for RT-DSBR is specific to the template. Fraction of repair products containing random insertions: (**A**) 2 bp, (**B**) 3 bp (excluding GAT), and (**C**) 4 bp insertions following Cas9-induced breaks repaired with either no donor or transcript RNA containing the GAT mutational signature, as measured by RT-DSBR (n>5). Statistical significance was assessed using an unpaired Student’s t-test. Error bars represent the standard error of the mean (± SEM). **D, E**, and **F** show the frequency of individual insertional signatures contributing to the overall values presented in **A, B**, and **C**, respectively (n>5). Error bars represent the standard error of the mean (± SEM).

**Extended Data Figure 6.**
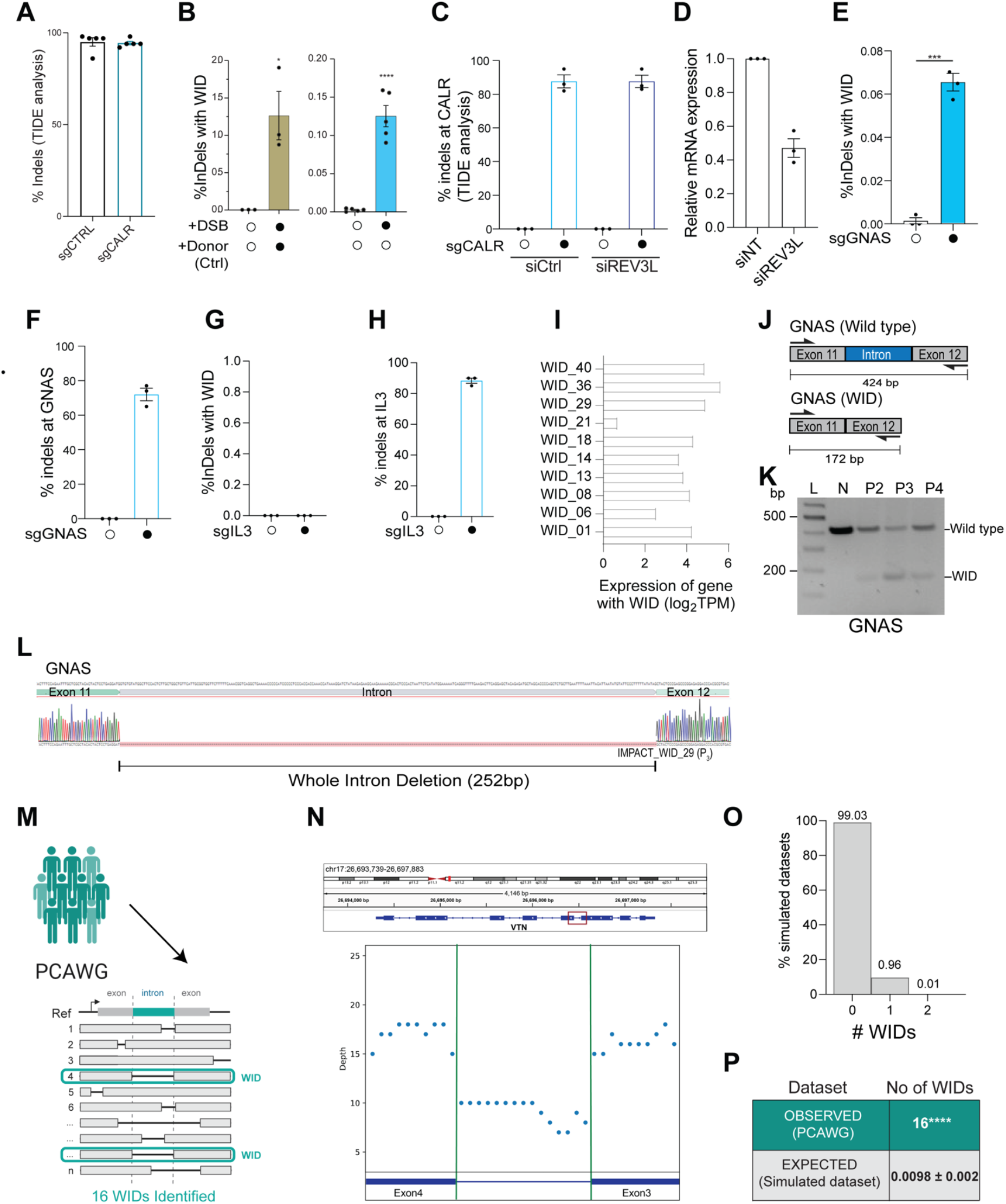
Repair via spliced RNA as a donor for RT-DSBR leads to WIDs. **A**, Editing efficiency for sgRNA targeting CALR in intron 2 calculated with TIDE analysis^70^ (n=3). **B**, Quantification of reads containing a precise deletion of the second intron at the CALR locus as a fraction of total reads after CRISPR-Cas9-induced break (n≥3). Left graph: WIDs were quantified in cells transfected with a donor that induced intron loss, compared to control cells without DSB induction. Right graph: A CRISPR-Cas9-mediated DSB was introduced without any exogenous donor template provided. WIDs driven by endogenous spliced mRNA were measured as a fraction of repair events. Statistical significance was assessed using unpaired Student’s t-test (* p < 0.05, ****p < 0.0001). Error bars represent the standard error of the mean (± SEM). **C**, Editing efficiency for sgRNA targeting CALR in intron two as depicted in (A). **D**, qRT-PCR to quantify the relative levels of REV3 transcripts in cells treated with control siRNA and siREV3L. **E**, Quantifying WIDs in cells treated with sgRNA targeting GNAS intron 11. **F**, TIDE analysis confirming efficient CRIPSR.Cas9 cleavage at the GNAS locus. **G**, Graph depicting the transcripts for CALR, GNAS, and IL3 obtained from human protein atlas for HEK293T cells. H, Analysis of intron loss upon cleaving the non-transcribed control gene, IL3. The left graph depicts WID events in cells treated with sgRNA targeting sgIL3. The right graph represented TIDE analysis confirming the cutting efficiency for sgRNA targeting IL3. I, Relative expression levels for the genes containing the WIDs in the tumor samples as determined by RNA-Seq analysis. **J**, Schematic of the exons spanning the WID in *GNAS* with the flanking primers used to confirm the sequence. A 172bp deletion was observed for the GNAS gene that maps precisely to the intron flanked by Exon 11-12. **K**, Agarose gel depicting the full-length band corresponding to the locus spanning Exon 11-12 in normal MCF-12A cells (N) and the shorted locus with the intron loss in the tumor sample in GNAS. P_2_-P_4_ represents three different patients from the MSK-IMPACT cohort. **L**, Sanger sequencing of the PCR products to confirm the presence of the WID in *GNAS*. **M**, Schematic of the pipeline to analyze the deletion profiles of PCAWG tumors. **N**, Example of a WID found in the VTN gene of a patient sample from the PCAWG cohort. **O**, 10,000 PCAWG-like cohorts were simulated, and the occurrence of WIDs was calculated. Graph representing the number of WID observed in the PCAWG-like simulated datasets. **P**, The number of expected WID was calculated after randomization of the deletion locations across the whole genome. Using Fisher’s exact test, empirical P-values were calculated by comparing the observed versus the 10,000 random values. (**** p < 0.0001).

## MATERIAL AND METHODS

### Cell culture

HEK293T cells were routinely grown with Dulbecco’s Modified Eagle’s Medium (DMEM) media supplemented with 10% bovine calf serum, 100 U/mL Penicillin-Streptomycin, 1% non-essential amino acids, 2mM L-glutamine. Cells were grown in a 37°C and 5% CO2 air incubator. Cells stably expressing the BFP reporter underwent selection with 20 ug/mL Blasticidin for ten days. Following selection with Blasticidin, cells were sorted with FACS. To generate POLQ KO clones, 10^6^ HEK293T-BFP cells were transfected with Cas9 expressing plasmid and two gRNAs targeting exon 3 of POLQ (Supplementary Table 1.5). Cells were enriched for Cas9 transfected cells. Individual colonies were seeded into a 96-well plate and grown to confluence before proceeding with genotyping. To generate TP53BP1 KO clones, 10^6^ HEK293T-BFP cells were transfected with a CRISPR-Cas9 ribonucleoprotein targeting exon 6 of TP53BP1 (Supplementary Table 1.5). Individual colonies were seeded into a 96-well plate and grown to confluence before proceeding with genotyping and western blotting. For AZT-treated samples, cells were treated for 24 hours with ten μM/mL of AZT (MilliporeSigma A2169)

### siRNA transfection

0.5×10^6^ cells were reverse transfected using RNAi max (Invitrogen) according to the manufacturer’s instructions with 30 pmol of siRNAs of the indicated genes (Dharmacon, Supplementary Table 1.3) or scrambled nontarget siRNA (Dharmacon, Supplementary Table 1.3).

### CRISPR-Cas9 Ribonucleoprotein preparation

The crRNA (IDT) and the tracrRNA (IDT) were resuspended to a final concentration of 100 mM in IDT buffer and mixed in an equimolar solution to a final concentration of 50 mM, heated at 95C for five minutes, and let cool at room temperature to form the crRNA:tracrRNA duplex. For each sample, 100 pmol of crRNA:tracrRNA duplex and 100 pmol of Cas9 enzyme (IDT) were diluted in PBS1X. The ribonucleoprotein formation reaction was incubated for 20 minutes at room temperature.

### Lentiviral production

For each transfection reaction, 5 ugs of RRE, 3 ugs of VSVG, and 2.5 ugs of REV plasmid DNA were mixed with 20.5 ugs of BFP plasmid DNA, 62 ug/mL polyethylenimine, and 150mM sodium chloride. The reaction was incubated at room temperature for 15 minutes and then added to a plate of 10 cm HEK293T cells at ~70-80% confluence. Cells were incubated at 37°C overnight. Fresh media was added, and the cells were left to recover for 6-8 hours before collecting the first viral supernatant. Fresh media was added, and after an additional 24 hours, the second viral supernatant was collected. This was repeated for the collection of a third viral supernatant.

### BFP-to-GFP conversion assay

To deliver Cas9 and the gRNA against the BFP sequence (Supplementary Table 1.5), a CRISPR-Cas9 ribonucleoprotein was formed. 10×6 cells were collected, washed with PBS1X, and resuspended in 100ul of SF nucleofection buffer (Lonza). 5 ul of the ribonucleoprotein mixture was added to the cells, as well as 1 ul of 100 μM of the repair donor. Reaction mixtures were electroporated in 4D Nucleocuvettes (Lonza) with the DS-150 program, incubated in the nucleocuvette at 37°C for 8 min with RPMI media, and transferred to culture dishes containing pre-warmed media. Cells were incubated for 72 hours and then analyzed for blue or green fluorescence via flow cytometry.

### Genomic DNA extraction and PCR amplification for BFP-to-GFP conversion Assay

To genotype cells into 96-well plates, cells were resuspended in gDNA “dirty” lysis buffer supplemented with 10mg/ml of Proteinase K. Cells were incubated overnight at 55C, and Proteinase K was inactivated by incubating the plate for five minutes at 65C. Genomic DNA extracted from BFP HEK293T cells was amplified via PCR with the primer pairs and 5 ml of extracted gDNA. The thermocycler was set for one denaturing cycle at 95C for three minutes, 35 cycles of denaturing at 95C for 15 seconds, annealing at 60C for 15 seconds, extension at 68C for 40 seconds, and one final extension cycle at 68C for five minutes before being held at 12C.

### Native PAGE

To check the purity of the chimera donors purchased from IDT, we run them on Native Polyacrylamide Gel Electrophoresis (Native PAGE) in the presence or absence of RNAse A. The separating gels were prepared at 6% from acrylamide and bis-acrylamide solutions 29:1 in TBE. Gels were pre-run at 160V for one hour before loading the samples. Before loading the chimera donors were treated with 10 ug of RNaseA for 30 minutes at 37C. Gels were run at 120 V until the ladder reached the end. Gels were stained with Ethidium bromide for 15 minutes and then washed three times in H2O. Typhoon GE Typhoon FLA 9000 Gel Scanner was used to detect the signal.

### Western blotting analysis

Cells were collected by trypsinization and lysed in RIPA buffer (25 mM Tris-HCl pH 7.6, 150 mM NaCl, 0.1% SDS, 1% NP-40, 1% sodium deoxycholate). After two cycles of water-bath sonication at medium settings, lysates were incubated at 4°C on a rotator for an additional 30 min. Lysates were clarified by centrifugation for 30 min at 14,800 rpm at 4°C, and the supernatant was quantified using the enhanced BCA protocol (Thermo Fisher Scientific, Pierce). Equivalent amounts of proteins were separated by SDS–PAGE and transferred to a nitrocellulose membrane. Membranes were blocked in 5% milk in TBST (137 mM NaCl, 2.7 mM KCl, 19 mM Tris-Base, and 0.1% Tween-20) or 5% BSA in TBST in the case of phosphorylated proteins for at least one hour at room temperature. Incubation with primary antibodies was performed overnight at 4°C. Membranes were washed and incubated with HRP-conjugated secondary antibodies at 1:5,000 dilution, developed with Clarity ECL (Biorad), and acquired with a ChemiDoc MP apparatus (Biorad) and ImageLab v.5.2. γ-tubulin was used as a loading control. The primary antibodies for Western blotting included FLAG (Clone M2, Sigma; 1: 10000 dilution) and γ-tubulin (GTU-88; Sigma Aldrich; 1: 5000 dilution). The secondary antibodies were mouse IgG HRP-linked (NA931, GE Healthcare; 1:5000) or rabbit IgG HRP-linked (NA934, GE Healthcare; 1:5000).

### qPCR validation of gene expression RT–qPCR

Total RNA was purified with the NucleoSpin RNA Clean-up (Macherey-Nagel) following the manufacturer’s instructions. Genomic DNA was eliminated by on-column digestion with DNase I. A total of 1 μg of RNA was reverse transcribed using iScript Reverse Transcription Supermix (Biorad), and cDNA was diluted 1:5 or 1:10. Reactions were run with ssoAdvanced SYBR Green Supermix (BioRad) with standard cycling conditions. Relative gene expression was normalized using *ACT1* as a housekeeping gene, and all calculations were performed using the ΔΔCt method. qPCR Primers are listed below in Supplementary Table 1.4.

### AAVS1-seq assay

0.15–0.25×10^6^ cells/well were seeded in six-well plates and treated with the respective siRNA as described above, 48 hours post-knock-down, cells were transfected with 2 μg of CRISPR plasmid (pX300) directed to the AAVS1 locus along with 10 μl of 10 μM donor oligo using Lipofectamine 3000 (Invitrogen). AAVS1 T2 CRISPR in pX330 was a gift from Masato Kanemaki (Addgene plasmid # 72833)^71^. 24 hours following transfection, cells were harvested and gDNA was extracted using the DNeasy Blood & Tissue Kit (Qiagen). To measure the use of transcript RNA as a template DSBR-seq, the pMJ1.19 plasmid transcribing a donor RNA complementary to the *AAVS1* locus was used. This contained the 3bp mutational signature interrupted by an artificial intron to help differentiate between the plasmid DNA and the transcribed RNA.

### DNA Library Preparation, HiSeq Sequencing for DSBR-seq

Initial DNA sample quality assessment, library preparation, and sequencing were conducted at Azenta (South Plainfield, NJ, USA). Genomic DNA samples were quantified using a Qubit 2.0 Fluorometer (Life Technologies, Carlsbad, CA, USA). Locus-specific primers (oMJ80 and oMJ81, Supplementary Table 1.6) were used to amplify target sequences. PCR products were cleaned up, and sequencing libraries were prepared using the NEBNext Ultra DNA Library Prep Kit according to the manufacturer’s protocol. In brief, amplicons were end-repaired and adenylated at the 3’ends. Adapters were ligated to the DNA fragments, and adapter-ligated DNA fragments were enriched and indexed with limited-cycle PCR. The adaptor-ligated sequencing libraries were validated on the Agilent TapeStation (Agilent Technologies, Palo Alto, CA, USA) and quantified by using Qubit 2.0 Fluorometer (Invitrogen, Carlsbad, CA) as well as by quantitative PCR (KAPA Biosystems, Wilmington, MA, USA). DNA libraries were multiplexed in equal molar mass and loaded on an Illumina HiSeq instrument according to the manufacturer’s instructions (Illumina, San Diego, CA, USA). Sequencing was performed using a 2×150 paired-end (PE) configuration; the HiSeq Control Software conducts image analysis and base calling on the HiSeq instrument. Illumina Reagent/kits for DNA library sequencing cluster generation and sequencing were used for enriched DNA sequencing.

Paired-end sequencing data were analyzed using CRISPResso, which aligns reads to the target region using a global alignment algorithm after merging read pairs with FLASh ^29^. Each unique mutational signature was identified utilizing a quantification window of 25 base pairs around the cut site (for sgRNA sequence, see Sup Table 1.5). For each sample, allele fractions for these events were calculated by counting the number of reads with respective mutational signatures identified by CRISPResso and dividing the count by the total reads. The fraction of indel reads was calculated by dividing the read count for mutational signatures by the number of reads harboring indels in the sample.

### Droplet digital PCR (ddPCR)

Custom assays specific for detecting mutations in AAVS1 were ordered through Bio-Rad. Primers and probes for ddPCR are listed below in Table 1.1. Cycling conditions were tested to ensure optimal annealing/extension temperature and optimal separation of positive from empty droplets.

After PicoGreen quantification, 9-27ng gDNA generated from the DSBR-seq assay was combined with locus-specific primers, FAM- and HEX-labeled probes, BamHI, and digital PCR Supermix for probes (no dUTP). All reactions were performed on a QX200 ddPCR system (Bio-Rad catalog # 1864001), and each sample was evaluated in technical replication of 2-8 wells. Reactions were partitioned into a median of ~14K droplets per well using the QX200 droplet generator. Emulsified PCRs were run on a 96-well thermal cycler using cycling conditions identified during the optimization step (95°C 10’; 40 cycles of 94°C 30’ and 60°C 1’; 98°C 10’; 4°C hold). Plates were read and analyzed with the QuantaSoft software to assess the number of droplets positive for each sample.

### Validation of the RNA-transcript-PCR/gel/sequencing

Total RNA was purified using the Quick-DNA/RNA™ Miniprep Plus (Zymo Research) following the manufacturer’s instructions. Genomic DNA was eliminated by on-column digestion with DNase I. A total of 2 μg of RNA was reverse transcribed using SuperScript™ IV VILO (Invitrogen). cDNA extracted from HEK293T cells transfected with the pMJ1.19 plasmid was amplified via PCR with either Primer Pair 1 (oMJ38-oMJ60) or Primer Pair 2 (oMJ39-oMJ61) (Supplementary Table 1.6) using Q5 master mix (NEB) under the following conditions: samples were denatured for 1 minute at 98°C for one cycle followed by 18 cycles of 98°C for 10 seconds, 55°C for 30 seconds, and 72°C for 20 seconds, the final extension step was performed for one cycle at 72C for 2 minutes. PCR samples were then run on a 1% Tris-acetate EDTA agarose gel and visualized using the Bio-Rad Chemidoc XRS system. The amplicons were confirmed using Sanger sequencing performed by Azenta (South Plainfield, NJ, USA).

### Genome-wide CRISPR/Cas9 screens

CRISPR screens were performed as previously described (Hart, 2015). HEK293T cells were transduced with a lentiviral DNA Damage Response library at a low MOI (~0.2–0.3) and selected with 4 ug/ml of puromycin for 48 h post-transduction, which was considered the initial time point (day 0). Cells were grown for 7 days and then divided into three subpopulations. One population was kept in culture for an additional 7 days and was considered the non-treated sample. Sample cell pellets were frozen at each time point for genomic DNA (gDNA) isolation. A library coverage of ≥500 cells per sgRNA was maintained at every step. gDNA from cell pellets was isolated using Midi Kit (ZymoResearch) and genome-integrated sgRNA sequences were amplified by PCR using the Q5 Mastermix (New England Biolabs Next UltraII). i5 and i7 multiplexing barcodes were added in a second round of PCR, and final products were sequenced on Illumina HiSeq2500 or NextSeq500 systems to determine sgRNA representation in each sample. MAGECK was used to identify essential genes ^39^.

### Tumor-data analysis (whole intron deletion identification)

Mutation data from the MSK-IMPACT solid tumor cohort (64,544 samples, 56,322 patients) was systematically scanned using a script to identify WIDs^51,53^. Canonical intron-exon boundaries were obtained from Ensembl transcript files (GRCh37). For the 73,030 deletions in the MSK-IMPACT cohort, intron-exon boundaries in the reference genome were compared with deletion boundaries to identify whole intron deletions. To account for alignment discrepancies, margins of +/-2bp were allowed between intron boundaries and deletion boundaries on both edges (Supplementary Tables 3.1).

Of the PCAWG cohort (1902 patients and samples), we identified whole intron deletions from 122,712 deletions longer than 10 base pairs using the same approach as mentioned above (Supplementary Tables 3.3).

### RNA extraction from tumor samples

Following Institutional Review Board (IRB) approval, formalin-fixed paraffin-embedded (FFPE) tissues of 10 cases were retrieved from the pathology archives of Memorial Sloan Kettering Cancer Center (MSK). Two pathologists (F.P. and T.V.) reviewed cases and included tumors arising from different anatomic locations (Supplementary Table 3.2). Cases were microdissected from ten eight-micron-thick histologic sections under a stereomicroscope (Olympus SZ61) to ensure a tumor content ≥80%. RNA was extracted using the RNAeasy FFPE kit (Qiagen) and subjected to RNA-sequencing at MSK Integrated Genomics Operation (IGO).

### RNA-seq on tumor samples

After RiboGreen quantification and quality control by Agilent BioAnalyzer, 0.5-1 µg of total RNA with DV200 percentages varying from 17-38% underwent ribosomal depletion and library preparation using the TruSeq Stranded Total RNA LT Kit (Illumina catalog # RS-122-1202) according to instructions provided by the manufacturer with 8 cycles of PCR. Samples were barcoded and run on a NovaSeq 6000 in a PE150 run, using the NovaSeq 6000 S4 Reagent Kit (300 Cycles) (Illumina).

### RNA-seq analysis

RNA sequencing reads were first examined using FASTQC^72^, then Illumina universal adapters were trimmed by cutadapt^73^. The trimmed reads were aligned to the GRCh37 human genome using STAR RNA-Seq aligner^74^, and then mapped single-end reads from transcripts were counted using the GenomicAlignments package in Bioconductor^75,76^. Read counts were further transformed into transcripts per million (TPM) normalized for gene length.

### Whole Intron Deletion PCR confirmation

Patient DNA samples were processed and procured from the MSKCC Integrated Genomics Operation core facility. Genomic DNA was amplified using primers, as mentioned in Supplementary Table 1.7 and Q5 master mix (NEB), under the following conditions. Samples were denatured for 3 minutes at 98°C for one cycle followed by 28 cycles of 98°C for 10 seconds, 60°C (GNAS) or 65°C (HLA) for 30 seconds, and 72°C for 20 seconds, the final extension step was performed for one cycle at 72°C for 2 minutes. PCR samples were then run on a 1% Tris-acetate EDTA agarose gel and visualized using the Bio-Rad Chemidoc XRS system. The amplicons were confirmed using Sanger sequencing performed by Azenta (South Plainfield, NJ, USA). The corresponding patients tested from the MSK-IMPACT cohort were: P1-IMPACT_WID_38, P2-IMPACT_WID_12, P3-IMPACT_WID_29, P4-IMPACT_WID_34

### Mathematical modeling

We developed a simulation strategy to quantify the likelihood of observing WIDs by random chance, rather than through any specific mechanism. This strategy simulates a cohort of deletions based on MSK-IMPACT data, taking into account both the genomic locations of mutations and the length of the deletion. We investigate whether the observed occurrence of WID in MSK-IMPACT data would exceed chance expectations based on simulated MSK-like cohorts.

The simulation approach learns the probability distribution of deletion lengths from the actual MSK-IMPACT data and uses this distribution as the probability to assign lengths to the simulated deletions. MSK-IMPACT is a targeted panel with only certain genomic regions being sequenced; here, we used these regions to reflect the space where simulated deletions could occur. Each interval within the MSK-IMPACT panel varies in its observed abundance of detected deletions in the actual MSK-IMPACT data, likely depending on interval length (some intervals are longer, and this could also increase the likelihood of deletions occurring) or even on biological reasons. To take this into account, we also used the probability distribution of deletions from the actual MSK-IMPACT data to assign probabilities to specific intervals in the simulation.

Each simulated cohort contains 73,030 deletions, mirroring the characteristics of the MSK-IMPACT cohort. Simulating a deletion involves three steps: 1) Randomly selecting a panel interval based on the observed probability distribution. 2) Randomly determining the starting position within the selected interval. 3) Randomly choose the deletion length according to the observed probability distribution. The end position of each simulated deletion is defined as the starting position plus the deletion length. Deletions must adhere to two constraints: staying within the same interval or ending at the start of the next interval to be detected by the MSK-IMPACT panel.

Each deletion was annotated after constructing a simulated dataset reflecting the MSK-IMPACT cohort, including simulated whole intron deletions. It should be noted that analyses based on the simulation strategy may be influenced by inherent randomness, leading to fluctuating results based on the random seed used. To address this, we create a cohort of 10,000 MSK-IMPACT-like deletion cohorts, employing different seed numbers for each cohort, ensuring distinct sets of simulated deletions. The p-value was calculated by determining the frequency at which a simulated cohort exhibits an equal or more significant number of WID compared to the actual MSK-IMPACT data.

Similarly, we generated 10,000 deletion cohorts resembling PCAWG data. Given that PCAWG samples were subjected to whole-genome sequencing (WGS), modifications to the simulation strategy were introduced. We focused the simulation on the subset of PCAWG deletions occurring within genes, as deletions in intergenic regions are irrelevant to this analysis. Within genes, the frequency of deletions based on the simulations was compared to the observed probability distribution in the actual PCAWG data. This approach sought to capture the relative abundance of deletions within genes, considering factors such as gene length.

### Intron loss-seq Assay

To deliver Cas9 and the gRNA against CALR intron 2, GNAS intron 11, and IL3 intron 4 (Supplementary Table 1.5), a CRISPR-Cas9 ribonucleoprotein was formed. 1X 10^6^ cells were collected, washed with PBS, and resuspended in 100ul of nucleofection buffer. SF buffer was used for HEK293T cells and SE buffer was used for PC9 cells (Lonza). 5 ul of the ribonucleoprotein mixture was added to cells transfected with either 1 ul of 100 μM of the repair donor (120bp ssDNA oligo chimera with 6 ribonucleotides spanning the exon-exon junction sequence (DNA^CALR-6R^) or no donor. Cells were electroporated using the 4D Nucleofector (Lonza) with the cell line specific program. Program DS-150 was used for HEK293T cells. Cells were incubated at 37°C for 8 min with RPMI media and then transferred to culture dishes containing pre-warmed media in which they were incubated for 72 hours. Genomic DNA was extracted using the Quick DNA Miniprep Kit (ZymoResearch) and then amplified by PCR with primers at the flanking exons (Supplementary Table 1.5) using Q5 Mastermix (New England Biolabs Next UltraII). i5 and i7 multiplexing barcodes were added in a second round of PCR, and final products were sequenced on Illumina HiSeq by the MSK IGO sequencing core using PE150 sequencing. CRISPResso2 was used to map reads to either the reference amplicon or the amplicon with a perfect intron deletion ^29^. A quantification window of 3 base pairs on either side of the exon-exon junction site was used to label and filter out imperfect intron loss as reads mapped to intron loss but containing insertions or deletions within this 6bp window. The percent of intron loss reads was calculated by dividing the read count for perfect intron loss by the total number of reads (reference + perfect intron loss + imperfect intron loss). For siRNA-mediated knockdown experiments, 1×10^6^ cells were seeded in 6 cm plates and treated with the respective siRNA as described above. 48 hours post-knock-down, cells were collected for nucleofection with CRISPR-Cas9 ribonucleoprotein.

### Statistics

All statistical analysis was performed with GraphPad Prism 9. Sample sizes and the statistical tests used are specified in the figure legends. Asterisks signify significance: * P< 0.05, **P< 0.01, ***P< 0.001, ****p < 0.0001.

